# The role of gene flow in rapid and repeated evolution of cave related traits in Mexican tetra, *Astyanax mexicanus*

**DOI:** 10.1101/335182

**Authors:** Adam Herman, Yaniv Brandvain, James Weagley, William R. Jeffery, Alex C. Keene, Thomas J. Y. Kono, Helena Bilandžija, Richard Borowsky, Luis Espinasa, Kelly O’Quin, Claudia P. Ornelas-García, Masato Yoshizawa, Brian Carlson, Ernesto Maldonado, Joshua B. Gross, Reed A. Cartwright, Nicolas Rohner, Wesley C. Warren, Suzanne E. McGaugh

## Abstract

Understanding the molecular basis of repeated evolved phenotypes can yield key insights into the evolutionary process. Quantifying the amount of gene flow between populations is especially important in interpreting mechanisms of repeated phenotypic evolution, and genomic analyses have revealed that admixture is more common between diverging lineages than previously thought. In this study, we resequenced and analyzed nearly 50 whole genomes of the Mexican tetra from three blind cave populations, two surface populations, and outgroup samples. We confirmed that cave populations are polyphyletic and two *Astyanax mexicanus* lineages are present in our dataset. The two lineages likely diverged ∼257k generations ago, which, assuming 1 generation per year, is substantially younger than previous mitochondrial estimates of 5-7mya. Divergence of cave populations from their phylogenetically closest surface population likely occurred between ∼161k - 191k generations ago. The favored demographic model for most population pairs accounts for divergence with secondary contact and heterogeneous gene flow across the genome, and we rigorously identified abundant gene flow between cave and surface fish, between caves, and between separate lineages of cave and surface fish. Therefore, the evolution of cave-related traits occurred more rapidly than previously thought, and trogolomorphic traits are maintained despite substantial gene flow with surface populations. After incorporating these new demographic estimates, our models support that selection may drive the evolution of cave-derived traits, as opposed to the classic hypothesis of disuse and drift. Finally, we show that a key QTL is enriched for genomic regions with very low divergence between caves, suggesting that regions important for cave phenotypes may be transferred between caves via gene flow. In sum, our study shows that shared evolutionary history via gene flow must be considered in studies of independent, repeated trait evolution.

## INTRODUCTION

Repeated adaptation to similar environments offers insight into the evolutionary process (Agrawal 2017; Gompel & Prud’homme 2009; Losos 2011; Rosenblum *et al.* 2014; Stern 2013; Stern & Orgogozo 2009). Predictable phenotypes are likely when repeated evolution occurs through standing genetic variation or gene flow, such that the alleles are identical by descent, and when populations experience shared, strong selection regimes (Rosenblum *et al.* 2014). Understanding repeated evolution, however, requires an understanding of the basic parameters of the evolutionary process, including how long populations have diverged, how many independent phenotypic origins have occurred, the extent of gene flow between populations, and the strength of selection needed to shape phenotypes (Roesti *et al.* 2014; Rosenblum *et al.* 2014; Rougemont *et al.* 2017; Stern 2013; Welch & Jiggins 2014).

The predictable phenotypic changes observed in cave animals offers one of the most exciting opportunities in which to study repeated evolution (Elmer & Meyer 2011). Cave animals also offer advantages over many systems in that the direction of phenotypic change is known (surface**→** cave) and coarse selection pressures are defined (e.g., darkness and low nutrient availability). The cavefish (*Astyanax mexicanus*) in northeastern Mexico have become a central model for understanding the evolution of diverse developmental, physiological, and behavioural traits (Keene *et al.* 2015). Many populations of *Astyanax mexicanus* exhibit a suite of traits common to other cave animals including reduced eyes and pigmentation (Borowsky 2015; Protas & Jeffery 2012). In addition, many populations possess behavioral and metabolic traits important for survival in dark, low-nutrient environments (Aspiras *et al.* 2015; Bibliowicz *et al.* 2013; Duboué *et al.* 2011; Jaggard *et al.* 2017; Jaggard *et al.* 2018; Jeffery 2001, 2009; Protas *et al.* 2008; Riddle *et al.* 2018; Salin *et al.* 2010; Varatharasan *et al.* 2009; Yamamoto *et al.* 2009; Yoshizawa *et al.* 2010). Over 30 populations of cavefish are documented (Espinasa *et al.* 2001; Mitchell *et al.* 1977), and conspecific surface populations allow a proxy for the ancestral conditions to understand repeated adaptation to the cave environment. Thus, *A. mexicanus* offers an excellent opportunity to evaluate the roles of history, migration, drift, selection and mutation to the repeated evolution of a convergent suite of phenotypes.

Despite evidence for the repeated evolutionary origin of cave-associated traits in *A. mexicanus*, the timing of cave invasions by surface lineages is uncertain, and the extent of genetic exchange between cavefish and surface fish is under debate (Bradic *et al.* 2012; Bradic *et al.* 2013; Coghill *et al.* 2014; Fumey *et al.* 2018; Gross 2012; Hausdorf *et al.* 2011; Ornelas-García *et al.* 2008; Porter *et al.* 2007; Strecker *et al.* 2003; Strecker *et al.* 2004; Strecker *et al.* 2012). High amounts of gene flow between cave and surface populations and among cave populations may complicate conclusions regarding repeated evolution (Ornelas-García & Pedraza-Lara 2015). For instance, cavefish populations are polyphyletic (Bradic *et al.* 2012; Bradic *et al.* 2013; Coghill *et al.* 2014; Gross 2012; Ornelas-García *et al.* 2008; Strecker *et al.* 2003; Strecker *et al.* 2012). In principle, this pattern is also consistent with a single invasion of the cave system, subterranean spread of fish, and substantial gene flow of individual caves with their nearest respective surface population (suggested by Coghill *et al.* 2014; Espinasa & Borowsky 2001). Alternatively, gene exchange among caves could result in a shared adaptive history even if cave populations were founded independently.

Here, we conduct an extensive examination of the gene flow between cave populations and between cave and surface populations of *A. mexicanus* to understand repeated evolutionary origins of cave-derived phenotypes and the potential for gene flow to enhance or impede adaptation to caves. Since we employ whole genome resequencing as opposed to reduced representation methods, we were able to calculate absolute measures of divergence (*d*_XY_), compare to relative measures (pairwise F_ST_) (reviewed in Cruickshank & Hahn 2014; Ellegren 2014; Lowry *et al.* 2016), and demonstrate that pairwise F_ST_ is predominantly driven by heterogeneous diversity across populations which obscured accurate inferences of divergence and gene flow among populations in past studies (Charlesworth 1998). Our analyses reveal gene flow between cave populations is much more extensive than previously appreciated, cave and surface populations regularly exchange genes (Wilkens & Strecker 2003), and surface and cave populations have diverged recently. We show that given these demographic parameters, repeated evolution of cave phenotypes may reflect the action, rather than the relaxation of natural section. Additionally, we present evidence that at least one region of the genome important for cave-derived phenotypes may be transferred between caves, suggesting that gene flow among caves may play a role in the maintenance and/or origin of cave phenotypes.

## METHODS

### Sampling, DNA extractions, and sequencing

We sampled five populations of *Astyanax mexicanus* cave and surface fish from the Sierra de Guatemala, Tamaulipas, and Sierra de El Abra region, San Luis Potosí, Mexico: Molino, Pachón, and Tinaja caves and the Rascón and Río Choy surface populations (Figure 1). Two main lineages of cave and surface populations are often referred to as ‘new’ and ‘old’ in reference to when the lineages reached northern Mexico, each with independently evolved cave populations. Populations in Molino cave and Río Choy are considered ‘new’ lineage fish, and the populations of Pachón cave and Tinaja cave are typically classified as ‘old’ lineage populations (Bradic *et al.* 2012; Coghill *et al.* 2014; Dowling *et al.* 2002; Ornelas-García *et al.* 2008).

**Figure 1.**
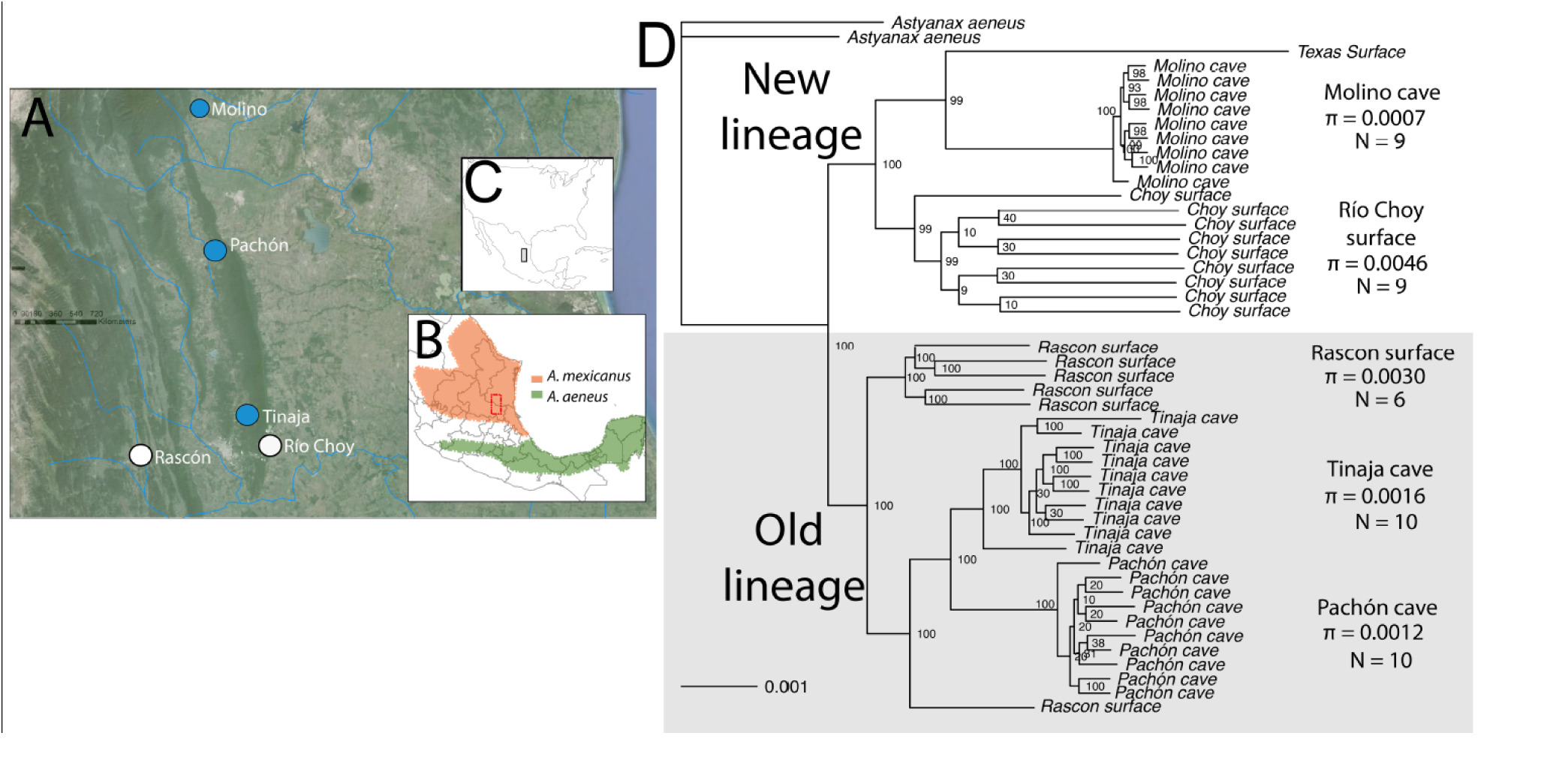
**A.** Map of caves (red), surface populations (blue), and additional unsampled cave populations (grey) **B.** Location of *A. mexicanus* range with sampled area in red box and *A. aeneus* range, **C.** Location in North and Central America of the focal sampled area. Map adapted from (Ornelas-García & Pedraza-Lara 2015). **D.** Phylogenetic inference from the largest scaffolds that comprised approximately 50% of the genome using a maximum likelihood tree search and 100 bootstrap replicates in RAxML v8.2.8(Stamatakis 2014). π is from Table 1. Old lineage and new lineage populations delineated by past studies are strongly supported. Branch-lengths correspond to phylogenetic distance and confirm cave populations (especially Molino cave) are less diverse than surface populations.

Rascón was selected because *cytochrome b* sequences are most similar to old lineage cave populations (Ornelas-García *et al.* 2008) and it is part of the river system of Rio Gallinas, which ends at the 105 m vertical waterfall of Tamul into Rio Santa Maria/Tampaon. It is hypothesized that new lineage surface fish could not overcome the 105 m waterfall and could not colonize the Gallinas river (e.g. Rascón population). Thus, Rascón likely has remained cut off from nearby surface streams inhabited by new lineage populations.

Fin clips were collected from fish in Spring 2013 from Pachón cave, Río Choy (surface), and a drainage ditch near the town of Rascón in San Luis Potosí, Mexico. Samples from Molino cave were collected by W. Jeffery, B. Hutchins, and M. Porter in 2005, and Pecos drainage in Texas in 1994 by W. Jeffery and C. Hubbs. Samples from Tinaja cave were collected in 2002 and 2009 by R. Borowsky.

We complemented our sequencing of these populations with three additional individuals. First, we sequenced an *A. mexicanus* sample from a Texas surface population. While this population is not of direct interest to this study, it has been the focus of many Quantitative Trait Loci (QTL) studies (e.g., O’Quin *et al.* 2015), and therefore it’s evolutionary relationship to our populations is of interest. Second, we aimed to sequence an outgroup to polarize mutations in *A. mexicanus*. To do so, we first sequenced two samples of a close congener *Astyanax aeneus* (Mitchell *et al.* 1977; Ornelas-García & Pedraza-Lara 2015) from Guerrero, Mexico. However, despite being separated from the sampled *Astyanax mexicanus* populations by the Trans-Mexican Volcanic Belt (Mitchell *et al.* 1977; Ornelas-García & Pedraza-Lara 2015), we found a complex history of gene flow between *Astyanax aeneus* and *Astyanax mexicanus*. We therefore sequenced a more distant outgroup, the white skirt tetra (long-finned) *Gymnocorymbus ternetzi*. *Gymnocorymbus ternetzi* was used only for polarizing 2D site frequency spectra.

DNA was extracted using the Genomic-Tip Tissue Midi kits and DNEasy Blood and Tissue kit (Qiagen). We performed whole genome resequencing with 100bp paired-end reads on an Illumina HiSeq2000 at The University of Minnesota Genomics Center. Samples were prepared for Illumina sequencing by individually barcoding each sample and processing with the Illumina TruSeq Nano DNA Sample Prep Kit using v3 reagents. Five barcoded samples were pooled and sequenced in two lanes. In total, 45 samples were sequenced across 18 lanes. We resequenced nine Pachón cavefish, ten Tinaja cavefish, nine Molino cavefish, six Rascón surface fish, nine Río Choy surface fish and two *Astyanax aeneus* samples. We also made use of the *Astyanax* genome project to add another sample from the Pachón population, and obtain genotype information for the reference sequence. For this reference sample, we used the first read (R1) from all of the 100bp paired-end reads that were sequenced using an Illumina HiSeq2000 and aligned to the reference genome (McGaugh *et al.* 2014). The Texas population and the white skirt tetra were barcoded, pooled, and sequenced across two lanes of 125bp paired-end reads from a HiSeq2500 in High Output run mode using v3 reagents.

For all raw sequences, we trimmed and cleaned sequence data with Trimmomatic v0.30 (Lohse *et al.* 2012) and cutadapt v1.2.1 (Martin 2011) using the adapters specific to the barcoded individual, allowing a quality score of 30 across a 6bp sliding window, and removing all reads < 30 nucleotides in length after processing. When one read of a pair failed QC, its mate was retained as a single-end read for alignment. Post-processing coverage statistics were generated by the Fastx toolkit v0.0.13 (http://hannonlab.cshl.edu/fastx_toolkit/) and ranged from 6.8 to 11.9 fold coverage (mean = 8.87 fold coverage; median = 8.65 fold coverage Table 1, S1, S2), assuming a genome size estimate of 1.19 Gb (McGaugh et al. 2014) for the Illumina Hi-Seq 2000 samples. For the Pachón reference genome sequence, which was excluded from the measures above, the coverage was 16.68 fold. All new Illumina sequence reads were submitted to NCBI’s Sequence Read Archive (Table S3). We expect some heterozygosity drop-out with this relatively low level of coverage, but we sought to balance number of samples and depth of coverage in a cost-effective manner. Heterozygosity drop out would lower our diversity estimates, and if directional (e.g., due to reference sequence bias) could increase divergence between populations and lower estimates of gene flow. As detailed below, our data estimate more recent divergence times and higher estimates of gene flow than past studies.

**Table 1.** Basic statistics for population level resequencing. Coverage statistics are for reads cleaned of adapters, filtered for quality, and aligned to the *Astyanax mexicanus* reference genome v1.0.2. π included all sites (invariant, SNPs) except highly repetitive masked regions of the genome (∼33% of the genome), indels, and 3 nt on either side of an indel were excluded in the analysis. Each site was required to have six or more individuals for π to be calculated and the values given are a genome-wide average. Two individuals of *Astyanax aeneus*, one Texas surface *Astyanax mexicanus*, and white long-finned skirt tetra (*Gymnocorymbus ternetzi*) were also sequenced.

| Population | N | aligned coverage<br>mean (range) | Mean $\pi$ | stdev |
| --- | --- | --- | --- | --- |
| Pachón - cave | 9 + reference | 9.94 (7.65, 17.74) | 0.0012 | 0.0002 |
| Tinaja - cave | 10 | 10.05 (7.27, 12.35) | 0.0016 | 0.0004 |
| Rascón - surface | 6 | 10.90 (8.63, 13.15) | 0.0030 | 0.0003 |
| Molino - cave | 9 | 9.39 (7.47, 12.32) | 0.0007 | 3e-04 |
| Río Choy- surface | 9 | 9.27 (8.38, 10.26) | 0.0046 | 0.0004 |

### Alignment to reference and variant calling

The *Astyanax mexicanus* genome v1.0.2 (McGaugh et al. 2014) was downloaded through NCBI genomes FTP. Alignments of Illumina data to the reference genome were generated with the BWA-mem algorithm (Li 2013) in *bwa-0.7.1* (Li & Durbin 2009, 2010). Both Genome Analysis Toolkit v3.3.0 (GATK) and Picard v1.83 (http://broadinstitute.github.io/picard/) were used for downstream manipulation of alignments, according to GATK Best Practices and forum discussions (Auwera *et al.* 2013; DePristo *et al.* 2011; McKenna *et al.* 2010). Alignments of paired-end and orphaned reads for each individual were sorted and merged using Picard. Duplicate reads that may have arisen during PCR were marked using Picard’s MarkDuplicates tool and filtered out of downstream analyses. GATK’s IndelRealigner and RealignerTargetCreator tools were then used to realign reads that may have been errantly mapped around indels (insertions/deletions). Additional details are in Supplemental Methods.

The HaplotypeCaller tool in GATK v3.3.0 was used to generate GVCFs of genotype likelihoods for each individual. The GenotypeGVCFs tool was used to generate a multi-sample variant call format (VCF) of raw variant calls for all samples. Hard filters were applied separately to SNPs and indels/mixed sites using the VariantFiltration and SelectVariants tools (Table S4). Filtering variants was performed to remove low confidence calls from the dataset. The filters removed calls that did not pass thresholds for base quality, depth of coverage, and other metrics of variant quality.

Alleles with 0.5 frequency appeared to be overrepresented in the site frequency spectra, and the depth of coverage for alleles with 0.5 frequency was greater than the coverage at other frequencies. Upon examination, every individual was heterozygous and these alleles occurred in small tracts throughout the genome. We concluded these were likely paralogous regions (teleost fish have an ancient genome duplication, Hoegg *et al.* 2004; Meyer & Van de Peer 2005). Molino was the most severely impacted population, likely due to Molino’s low diversity allowing for collapsing of paralogs (see below). To be conservative, we identified sites where 100% of individuals in any population were heterozygotes and excluded these sites in all analyses.

We generated a VCF file with variant and invariant sites, and subset it to include biallelic only SNPs where applicable. We excluded sites that were defined as repetitive regions from a Repeat Modeler analysis in McGaugh et al. (2014). Next, we removed sites that had less than six individuals genotyped in all populations. We also scanned for indels in the VCF file and removed the indel as well as 3 bp +/- around the indel. In total, 171,841,976 bp were masked from downstream analyses. Lastly, all sites that were tri or tetra-allelic were removed. Thus, sites were either invariant or biallelic. To reduce issues of non-independence induced by linkage disequilibrium, we analyzed a thinned set of biallelic SNPs by retaining a single SNP in each non-overlapping window of 150 SNPs for the ADMIXTURE, TreeMix, and *F*_*3*_ and *F*_*4*_ analyses, resulting in approximately 119,000 SNPs for those analyses.

### Population phylogeny and divergence

We generated a phylogenetic reconstruction to understand population relationships using 47 sampled individuals (excluding white skirt tetra, but including *A. aeneus* and Texas *A. mexicanus*). The population phylogeny estimation was implemented in RAxML v8.2.8 with the two *A. aeneus* specified as the outgroups. We converted the VCF to fasta format alignments by passing through the mvf format using MVFtools v3 (Pease & Rosenzweig 2015). This was done to preserve ambiguity codes at sites polymorphic in individuals. Sites with greater than 40% missing data were removed using TrimAl v1.2 (Capella-Gutiérrez *et al.* 2009). We performed 100 rapid bootstrapping replicates followed by an ML search from two separate parsimony trees. The ML tree with bootstrap support was drawn using the ape package v3.5 (Paradis *et al.* 2004) in R v3.2.1 (Team 2014) using the *A. aeneus* samples as the outgroup, however, our results are consistent even if the outgroup is left unspecified and this decision is justified by the observation that *A. aeneus* is more distant from all *A. mexicanus* populations than any are to one another, as measured by the mean number of pairwise sequence differences.

#### Intra- and inter-population average pairwise nucleotide differences

Average pairwise nucleotide diversity values (π) reflect the history of coalescence events within and among populations (Hudson 1990; Nei 1987) and are directly informative about the history and relationships of a sample. Here, we use the following notation for inter- and intrapopulation average pairwise nucleotide diversity values: intrapopulation values are designated with π, while interpopulation values are designated with *d*_XY_ where X and Y are different populations (e.g., *d*_Rascón-Pachón_). Pairwise nucleotide diversity estimates for fourfold degenerate sites were calculated for 46 individuals (excluding Texas *A. mexicanus* and white skirt tetra) from unphased genomic data using the VCF file with invariant and variant sites unless otherwise noted. Additional details for each analysis (e.g., window size, site class used) are given supplemental methods. We also performed windowed estimates along the entire genome to understand the fine-scale apportionment of ancestry.

We use these levels of diversity and divergence to estimate the difference in coalescent times between and within populations. In the absence of gene flow, this excess between populations can be translated into an estimate of divergence time (*d*_XY_ - π_anc_ = 2μT (Brandvain *et al.* 2014; Hudson *et al.* 1987), where T is the population split time, and using a diversity in surface population as a stand in for π_anc_). After removing early generation hybrids identified by ADMIXTURE, we use this approach to estimate this excess coalescent time between pairs of populations. However, given extensive evidence for gene exchange (below), this estimate will be more recent than the true divergence time, which we estimate with model-based approaches below.

In this and all other estimates of divergence time, below, we assume a neutral mutation rate of 3.5 × 10^−9^/bp/generation which is estimated using parent-offspring trios in cichlids (Malinsky *et al.* 2017). Though generation time in the wild has not been measured, in the laboratory, *Astyanax* is sexually mature as early as six months under optimal conditions, though this varies across laboratories (Borowsky 2008a; Jeffery 2001). Since conditions in the wild are rarely as optimal as in the laboratory, we assume generation time is one year, though, our general conclusions are consistent if generation time is only six months.

### Genome-wide tests for gene flow

#### ADMIXTURE

We estimated admixture (cluster membership) proportions for the individuals comprising the five populations studied, as well as *A. aeneus* samples, using ADMIXTURE v1.3.0 (Alexander & Lange 2011; Alexander *et al.* 2009). We conducted 10 independent runs for each value of K (the number of ancestral population clusters) from K = 2 to K = 6 and present results for the run with the K = 6 (i.e., the number of sampled biological populations).

#### F_*3*_ and F_*4*_ statistics

After removing early generation hybrids, we used *F*_*3*_ and *F*_*4*_-statistics (as implemented in TreeMix v1.12 (Pickrell & Pritchard 2012)) to test for historical gene flow. For computational feasibility, we calculated a standard error for each statistic using a block jackknife with blocks of 500 SNPs, assessing significance by a Z-score from the ratio of *F*_*4*_ to its standard error.

The *F*_*3*_ statistic (Peter 2016; Reich *et al.* 2009), represented as *F*_*3*_ (X;A,B), is calculated as E[(*p*_*X*_ - *p*_*A*_)(*p*_*X*_ - *p*_*B*_)], where X is the population being tested for admixture, A and B are treated as the source populations, and *p*_*X*_, *p*_*A*_, *p*_*B*_ are the allele frequencies. Without introgression, *F*_*3,*_ is positive, and a negative value occurs when this expectation is overwhelmed by a non-treelike history of three populations. Importantly, complex histories of mixture in populations A and/or B do not result in significantly negative *F*_*3*_ values (Patterson *et al.* 2012; Reich *et al.* 2009).

*F*_*4*_ tests assess ‘treeness’ among a quartet of populations (Reich *et al.* 2009). *F*_*4*_(P1,P2;P3,P4) is calculated as E[(*p*_*1*_ - *p*_*2*_)(*p*_*3*_ - *p*_*4*_)], and serves as a powerful test of introgression when P1, P2 and P3, P4 are sisters on an unrooted tree. A significantly positive *F*_*4*_ value means that P1 and P3 are more similar to one another than expected under a non-reticulate tree — a result consistent with introgression between P1 and P3, or between P2 and P4. Alternatively, a significantly negative *F*_*4*_ value is consistent with introgression between P1 and P4 or between P2 and P3. Regardless of the sign of this test, F4 values may reflect introgression events involving sampled, unsampled, or even extinct populations.

#### TreeMix

We used the Treemix program v1.12 (Pickrell & Pritchard 2012) to visualize all migration events after removing early generation hybrids. Rather than representing population relationships as a bifurcating tree, TreeMix models population relationships as a graph in which lineages can be connected by migration edges. We used the same thinned biallelic SNPs as above. As with *F*_*3*_ and *F*_*4*_, we removed early generation hybrids from this analysis. We first inferred the maximum-likelihood tree and then successively added single migration events until the proportion of variance explained by the model plateaued (Pickrell & Pritchard 2012).

### Demographic modeling

We modeled the demographic history of pairs of populations to further elucidate their relationships and migration rates. To conduct demographic modeling, we generated unfolded joint site frequency spectra (2D SFS) from the invariant sites VCF using a custom Python script. Detailed explanation of parameters and a user-friendly guide is available here: https://github.com/TomJKono/CaveFish_Demography/wiki

White skirt tetra (*Gymnocorymbus ternetzi*) was used as the estimated ancestral state for all 10 pairwise combinations of Río Choy surface, Molino cave, Pachón cave, Rascón surface, and Tinaja cave. Recently admixed individuals were excluded from the comparisons. To reduce the effect of paralogous alignment from the ancient teleost genome duplication, sites at 50% frequency in both populations were masked from the demographic modeling analysis. Sites were excluded if they were heterozygous in the representative outgroup sample or had any missing information in either of the populations being analyzed. In addition, sites were excluded due to indels or repetitive regions as described above. Aside from these exclusions, all other sites, including invariant sites were included in all 2D SFS and the length of these sites make up locus length.

A total of seven demographic models were fit to each 2D SFS to infer the timing of population divergence, effective migration rates, and effective population sizes using ∂a∂i 1.7.0 (Gutenkunst *et al.* 2009). The seven models were derived from previously published demographic modeling analysis (Tine *et al.* 2014). Briefly, the seven models are as follows: SI - strict isolation, SC - secondary contact, IM - isolation with migration, AM - ancestral migration, SC2M - secondary contact with two migration rates, IM2M - isolation with migration with two migration rates, and AM2M - ancestral migration with two migration rates (see Figure S9 of Tine *et al.* 2014). Two migration rates within the genome appeared to fit the data better in previous work (Tine *et al.* 2014), as this approach allows for heterogeneous genomic divergence (M which is most often the largest migration rate within the genome, Mi which is usually the lowest migration rate within the genome). Descriptions of the parameters and parameter starting values are given in Table S5 and Supplementary Methods. For each replicate, Akaike Information Criterion (AIC) values for each model were converted into Akaike weights. The model with the highest mean Akaike weight across all 50 replicates was chosen as the best-fitting model (Rougeux *et al.* 2017; Table S6; similar to Wagenmakers & Farrell 2004).

The best-fitting demographic model for each 2D SFS comparison was used to generate estimates of the population parameters. Scaling from estimated parameters to real-value parameters was done assuming a mutation rate (μ) of 3.5 × 10^−9^ per bp per generation (Malinsky *et al.* 2017) and a locus length (L) of the number of sites used to generate the 2D SFS. For effective population size estimates, we estimated the reference population size, Nref, as the mean “theta” value from all 50 replicates of the best-fitting model multiplied by 1/(4μL). The effective population sizes of the study populations were then estimated as a scaling of Nref. To estimate the per-generation proportion of migrants, we scaled the mean total migration rates from ∂a∂i by 1/(2Nref). Total times since divergence for pairs of populations were estimated as the sum of the split time (“Ts”) and time since secondary contact commenced (“Tsc”) estimates for each population pair.

### Modeling of selection needed for cave alleles to reach high frequency

With demographic estimates provided by our whole genome sequencing and ∂a∂i, we estimated the selection coefficients needed to bring alleles associated with the cave phenotype to high frequency. We implemented the 12-locus additive alleles model by Cartwright *et al.* (2017) with the parameters estimated for Molino cave, as this is the population where selection would need to be strongest to overcome the effects of drift. We estimated the one-way mutation rate of loci (μ) to be on the order of 1 × 10^−6^ (c.f. 3.5 × 10^−9^/bp/generation * 1170bp, which is the median gene length across the genome). Other parameters were N = 7,335 (Molino Ne), h = 0.5 (additive alleles), k = 12 (number of loci), and Q = 0.1 (surface allele frequency of cave-favored alleles). The simulation model was adapted from Cartwright *et al.* (2017) with the addition that the cave population was isolated for a period of time before becoming connected again with the surface. Consistent with our demographic models for Molino and Río Choy, migration rates spanned between 10^−6^ – 10^−5^ with 91,000 generations of isolation followed by 71,000 generations of connection. We also conducted simulations with the parameters from Tinaja cave as this is the cave population where selection would need to be weakest to overcome the effects of drift. All parameters were the same as above except N = 30,522 (Tinaja Ne), and 170,000 generations of isolation were followed by 20,000 generations of connection.

### Candidate regions introgressed between caves

To identify regions of the genome that were likely transferred between cave populations or between cave and surface populations and were linked to cave-associated phenotypes, we implemented an outlier approach incorporating pairwise sequence differences and F_ST_. To determine potential candidate regions, we utilized 5kb *d*_XY_ windows using all biallelic sites, not just four-fold degenerate, since so few sites would be available. We first phased the biallelic variant sites for all samples using BEAGLE (version 4.1; Browning & Browning 2007). We then determined the number of invariant sites (from the full genome VCF) and variant sites in each 5kb window to calculate average pairwise diversity. We then identified 5kb regions in which *d*_XY_ was in the lower 5% tail of the genomic distribution of *d*_XY_ for all three pairwise combinations of cave populations.

We identified genes with low *d*_XY_ across all caves (lowest 5% of *d*_XY_ values genome-wide across all three pairwise comparisons) but substantial divergence with either surface population as measured by F_ST_ or *d*_XY_. For F_ST_ outliers, we required that π per gene for both surface fish populations must be greater than the lowest 500 π values for genes across the genome. This requirement protects, in part, against including regions that are low diversity across all populations due to a feature of the genome.

To put the results in context of phenotypes mapped to the genome, we created a database of prior QTL from several key studies (Kowalko *et al.* 2013a; Kowalko *et al.* 2013b; O’Quin *et al.* 2013; Protas *et al.* 2007; Protas *et al.* 2008; Yoshizawa *et al.* 2015; Yoshizawa *et al.* 2012b) and used overlapping markers between studies to position QTL relative to the linkage map in O’Quin *et al.* (2013). For markers in (Kowalko *et al.* 2013b), we used Blastn to place the marker on a genomic scaffold and placed the QTL from (Kowalko *et al.* 2013b) in our database as locating to the entire scaffold. Our QTL database is given in Table S7.

## RESULTS

### Population phylogenetic reconstruction

We inferred the population phylogeny using half the nuclear genome (invariant and variant sites, but no mitochondrial sites) in the program RAxML. Our phylogenetic tree clearly demarcates two lineages and indicates that the Rascón surface population and the Tinaja and Pachón cave populations form a monophyletic clade (Figure 1; often referred to as ‘old lineage’). Similarly, the Río Choy surface and Molino cave populations form a monophyletic clade (referred to as ‘new lineage’). Thus, this phylogeny indicates that many cavefish traits may be polyphyletic (i.e., evolved through repeated evolution).

Branch lengths illustrate the low diversity in cave populations as compared to surface populations. In agreement with a previous study, diversity is lower in the Molino cave population than other populations tested (Bradic *et al.* 2013). While all populations and the two lineages have high bootstrap support (≥ 99), bootstrap support with genomic-scale data provides little information about the evolutionary processes or the distribution of alternative topologies across the genome (Yang & Rannala 2012), thus, we use a series of tests below to explore this further.

### Diversity and divergence

Patterns of diversity within populations, π, and interpopulation divergence, *d*_XY_, provide a broad summary of the coalescent history within and between populations. We present pairwise sequence differences at fourfold degenerate sites between all pairs of individuals (Figure 2). The ‘striped’ individuals in Figure 2B correspond to early generation hybrids as inferred by ADMIXTURE (Figure 2A). We removed these putative early generation hybrids from our summary of diversity within and divergence between populations presented in Table 2.

**Figure 2.**
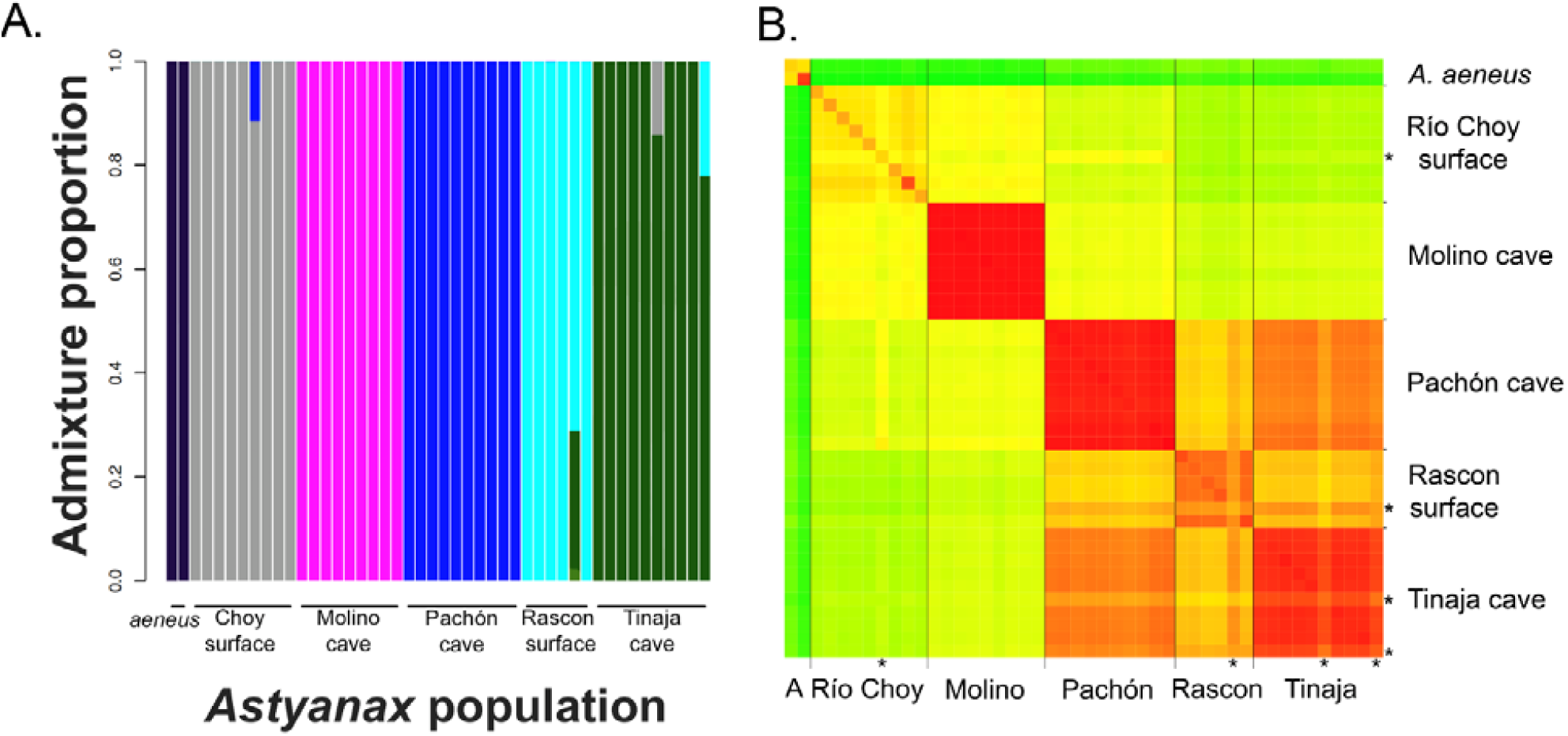
A. ADMIXTURE analysis suggests contemporary gene flow between old and new lineages and cave and surface populations. Analysis was performed on the thinned Biallelic SNP VCF to account for linkage disequilibrium between SNPs. We were interested in the relationships among the five nominal populations and the putative outgroup rather than an unknown, best-supported number of clusters, thus, we set the clusters to six. Additional analyses performed with other cluster numbers are given in the supplementary material. **B.** Pairwise nucleotide similarity (e.g., *d*_XY_, π) between all sampled individuals for fourfold degenerate sites genome-wide. Green = less similar, Red = more similar. Particular individuals potentially exhibiting admixture are indicated with an asterisk (e.g., Rascón 6, Tinaja 6, Río Choy 14, Tinaja E [bottom right corner]). *Astyanax aeneus* is represented by two total samples and indicated by the A on the x-axis.

**Table 2.** Inter- and intra-population average pairwise nucleotide diversity for fourfold degenerate sites (*d*_XY_ and π, respectively) and estimated population split times for the six populations sampled. Values in italics on the diagonal are the estimates of mean π for that population, values below the diagonal are *d*_XY_ for the population pair. Values above the diagonal are estimates, in generations, of the population split times (see Methods). In this table, four early-generation hybrids were excluded (e.g., Rascón 6, Tinaja 6, Choy 14, Tinaja E). Resulting in the sample sizes: *A. aeneus*: N = 2; Río Choy: N = 8; Molino: N = 9; Rascón: N = 5; Pachón: N = 9; Tinaja: N = 8 in the final calculations. N/A means the divergence time was negative. Divergence time does not take into account gene flow, therefore, is likely underestimated.

|  | *A. aeneus* | Río Choy | Molino | Pachón | Rascón | Tinaja |
| --- | --- | --- | --- | --- | --- | --- |
| *A. aeneus* | 0.00297 | 208,571 | 208,571 | 164,286 | 144,286 | 158,571 |
| Río Choy | 0.00443 | 0.00300 | 50,000 | 81,429 | 110,000 | 108,571 |
| Molino | 0.00443 | 0.00332 | 0.000738 | 61,571 | 84,857 | 79,714 |
| Pachón | 0.00412 | 0.00354 | 0.003401 | 0.001000 | 857 | N/A |
| Rascón | 0.00398 | 0.00374 | 0.003564 | 0.002976 | 0.002067 | N/A |
| Tinaja | 0.00408 | 0.00373 | 0.003528 | 0.002251 | 0.002854 | 0.001291 |

Genome-wide average diversity within cave populations (π_4fold degen_: Molino = 0.00074, Tinaja = 0.00129, Pachón = 0.00100; Table 2) is substantially lower than diversity within surface populations (π_4fold degen_: Río Choy = 0.00300, Rascón = 0.00207), reflecting a decrease in effective population size in caves (sensu Avise & Selander 1972). Molino cave is the most homogenous. These results are consistent with the short branches in cave populations observed in the phylogenetic tree produced by RAxML (Figure 1). In contrast, the surface population Río Choy is the most diverse in our sample — so much so that two Río Choy fish may be more divergent than fish compared between any of the old lineage populations (e.g., Tinaja and Rascón).

Genome-wide average divergence between two lineages (*d*_XY 4fold degen_ = 0.00340 - 0.00374, Table 2) exceeds divergence between cave and surface populations within lineages (*d*_XY 4fold degen_ Molino-Río Choy = 0.00332; Tinaja-Rascón = 0.00285; Pachón-Rascón = 0.00298; Table 2). Thus, old and new lineages diverged prior to (or have experienced less genetic exchange than) any of the cave-surface population pairs. The two old lineage caves are the least diverged populations in our sample (*d*_XY 4fold degen_: Pachón-Tinaja = 0.00225). These results are consistent with both the observation of monophyly of old and new lineages and the observation that old lineage cave populations are sister taxa in the RAxML tree (Figure 1).

Divergence between populations suggests that a simple bifurcating tree does not fully capture the history of these populations. For example, divergence between the new lineage cave population (Molino) and the geographically closer old lineage cave population (Pachón) is less than divergence between Molino and the geographically further old lineage populations (Tinaja and Rascón; Table 2). Likewise, all old lineage populations are closer to Molino cave than they are to Río Choy. These results suggest gene flow between Molino cave and the old lineage populations, which is supported by additional analyses.

### Genome-wide tests suggest substantial historical and contemporary gene flow

#### ADMIXTURE

We focus our discussion on K = 6, the case where the number of clusters matches our presumed number of populations (Figure 2A), and additional cluster sizes are presented in (Figure S1). While, individuals largely cluster exclusively with others from their sampled population, we also observe contemporary gene flow between cave and surface populations, as well between new and old lineage taxa (Figure 2A, S1).

Specifically, there appears to be reciprocal gene exchange between the Tinaja cave and the old lineage surface population (a Tinaja individual shares 22% cluster membership with Rascon surface and a Rascon individual shares 29% membership with Tinaja; Figure 2A, S1, S2). Another Tinaja sample appears to be an early generation hybrid with the new lineage surface population (a Tinaja individual with 14% membership with Río Choy). One new lineage surface sample appears to have recent shared ancestry with samples from Pachón (i.e., Río Choy individual with 12% cluster membership with Pachón).

Although unsupervised clustering algorithms may show signs of admixture even when none has occurred (Falush *et al.* 2016), corroboration of ADMIXTURE results with supporting patterns of pairwise sequence divergence (Figure 2, S1, S2) in putatively admixed samples (Rascón 6, Tinaja 6, Choy 14, Tinaja E) strongly suggests that early generation hybrids between cave and surface populations and old and new lineages are present in current-day populations.

While the existence of early generation hybrids is consistent with long-term genetic exchange, such hybrids do not provide evidence for historical introgression. We, therefore, rigorously characterize the history of gene flow in this group, while removing putative recent hybrids identified from ADMIXTURE and pairwise sequence divergence to ensure that our historical claims are not driven by a few recent events.

#### F_3_ and F_4_ statistics

To test for historic genetic exchange, we calculated *F*_*3*_ and *F*_*4*_ “tree imbalance tests,” after removing early generation hybrids identified by ADMIXTURE. Throughout the section below, gene flow is supported by *F*_*4*_ statistics between the two underlined taxa and/or the two non-underlined taxa in the four-taxa tree. Further, we interpret the *F*_*4*_ statistics in relation to the ranking of the *F*_*4*_ scores. Extreme *F*_*4*_ scores may be the result of both pairs (e.g., A-C and B-D) exchanging genes, amplifying the *F*_*4*_ score beyond what the score would be if gene flow occurred only between a single pair within the quartet.

*F*_*3*_ and *F*_*4*_ tests convincingly show that Rascón, the old lineage surface population, has experienced gene flow with a lineage more closely related to *A. aeneus* than to the other individuals in our sample (Table 3). This claim is supported by numerous observations including the observations that, while all *F*_*3*_ tests including Rascón are significantly negative (i.e., supporting admixture), the most extreme *F*_*3*_ statistic is from a test with Rascón as the target population and *A. aeneus* and Tinaja as admixture sources, reflecting that both *A. aeneus* and Tinaja have likely hybridized extensively with Rascón. Further, as measured by *d*_XY_, Rascón is more closely related to *A. aeneus* than other populations (Table 2). Additionally, all *F*_*4*_ tests of the form (*A. aeneus*, new lineage; old lineage cave, Rascón) are significantly negative, meaning that Rascón is genetically closer to *A. aeneus* than expected given a simple tree (Figure S3). Therefore, *A. aeneus* is an imperfect outgroup to *A. mexicanus*.

**Table 3.** *F*_3_ statistics for significant configurations out of all possible three population configurations (X;A,B) where X is the population tested for admixture. A significantly negative *F*_3_ statistic indicates admixture from populations related to A and B. Notably, we conducted the *F*_3_ statistics without individuals showing recent evidence of admixture. Sample sizes were as follows: *A. aeneus*: N = 2; Río Choy: N = 8; Molino N = 9; Rascón N = 5; Pachón N = 9; Tinaja N = 8. Any z-score below -1.645 passes the critical value for significance at α = 0.05. Tests are ordered most-least extreme z-scores. All other confirmations were not significant for introgression.

| (X; A, B) | $F_3$-statistic | standard error | z-score |
| --- | --- | --- | --- |
| Rascón;*A. aeneus*,Tinaja | -0.000925258 | 5.63403e-05 | -16.4227 |
| Rascón;Río Choy,Tinaja | -0.000733913 | 4.96641e-05 | -14.7775 |
| Rascón;Molino,Tinaja | -0.000720705 | 5.1705e-05 | -13.9388 |
| Rascón;*A. aeneus*,Pachón | -0.000552022 | 6.04747e-05 | -9.12813 |
| Rascón;*A. aeneus*,Molino | -0.000373176 | 6.53244e-05 | -5.71266 |
| Rascón;Río Choy,Pachón | -0.000204061 | 6.02999e-05 | -3.3841 |
| Rascón;Molino,Pachón | -0.000146049 | 6.18243e-05 | -2.36232 |
| Río Choy; *A. aeneus*,Molino | -0.000101532 | 5.50833e-05 | -1.84324 |

Our results also show that the new lineage surface population, Río Choy, experienced gene flow with an unsampled outgroup. A significantly negative *F*_*3*_ value demonstrates Río Choy (the new lineage surface population) is an admixture target with Molino (new lineage cave) and *A. aeneus* as admixture sources (Table 3). This claim is bolstered by significantly positive *F*_*4*_ values in all comparisons of the form (*A. aeneus*, old lineage; Río Choy, Molino) in Figure S3. However, because *d*_XY_ between *A. aeneus* and both new lineage populations are equivalent, we suggest this is an example of “the outgroup case” for *F*_*3*_ (Patterson *et al.* 2012) in which admixture is attributed to an uninvolved outgroup (in this case, *A. aeneus*) rather than the unsampled admixture source.

We uncover evidence for admixture between the old lineage cave population, Pachón, and both new lineage populations (Molino cave and Río Choy surface), as observed in early generation hybrids in ADMIXTURE. Specifically, the significantly negative *F*_*4*_ values for (*A. aeneus*, new lineage; Pachón, Tinaja) (Figure S3) are consistent with gene flow between both new lineage populations and the old lineage Pachón cave, and/or gene flow between Tinaja and *A. aeneus*. Because no other evidence suggests gene flow between Tinaja and *A. aeneus*, we argue that this result reflects gene flow between Pachón and the new lineage populations. This claim is further supported by the observation that *d*_XY_ between new lineage populations and Pachón is consistently lower than *d*_XY_ between new lineage populations and the other old lineage populations (Table 2).

Additionally, our analyses are consistent with gene flow between the old lineage Tinaja cave and the old lineage surface population, Rascón. Again, this result complements the discovery of two early generation hybrids between these populations in our ADMIXTURE analysis. This claim is supported by the significantly positive *F*_*4*_ value for (*A. aeneus*, Rascón; Pachón, Tinaja). Since we lack evidence of Pachón - *A. aeneus* admixture, this significant *F*_*4*_ statistic is likely driven by Rascón - Tinaja admixture.

Despite gene flow among most population pairs, there is one case in which tree imbalance tests failed to reject the null hypothesis of a bifurcating tree: *F*_*4*_ tests of the form (new lineage, new lineage; old lineage, old lineage) (Figure S3). This result is counter to both our observation of early generation hybrids (between the new and old lineages observed in ADMIXTURE), and the rampant gene flow observed in other *F*_*4*_ - statistics, as well as TreeMix (Figures 3, S4), *d*_XY_ nearest neighbor proportions (Figures S5-S7) and Phylonet (Figure S8). In these cases, it is also plausible that *F*_*4*_ scores that do not differ from zero may reflect opposing admixture events that cancel out a genome-wide signal, rather than an absence of introgression (see Reich *et al.* 2009).

**Figure 3.**
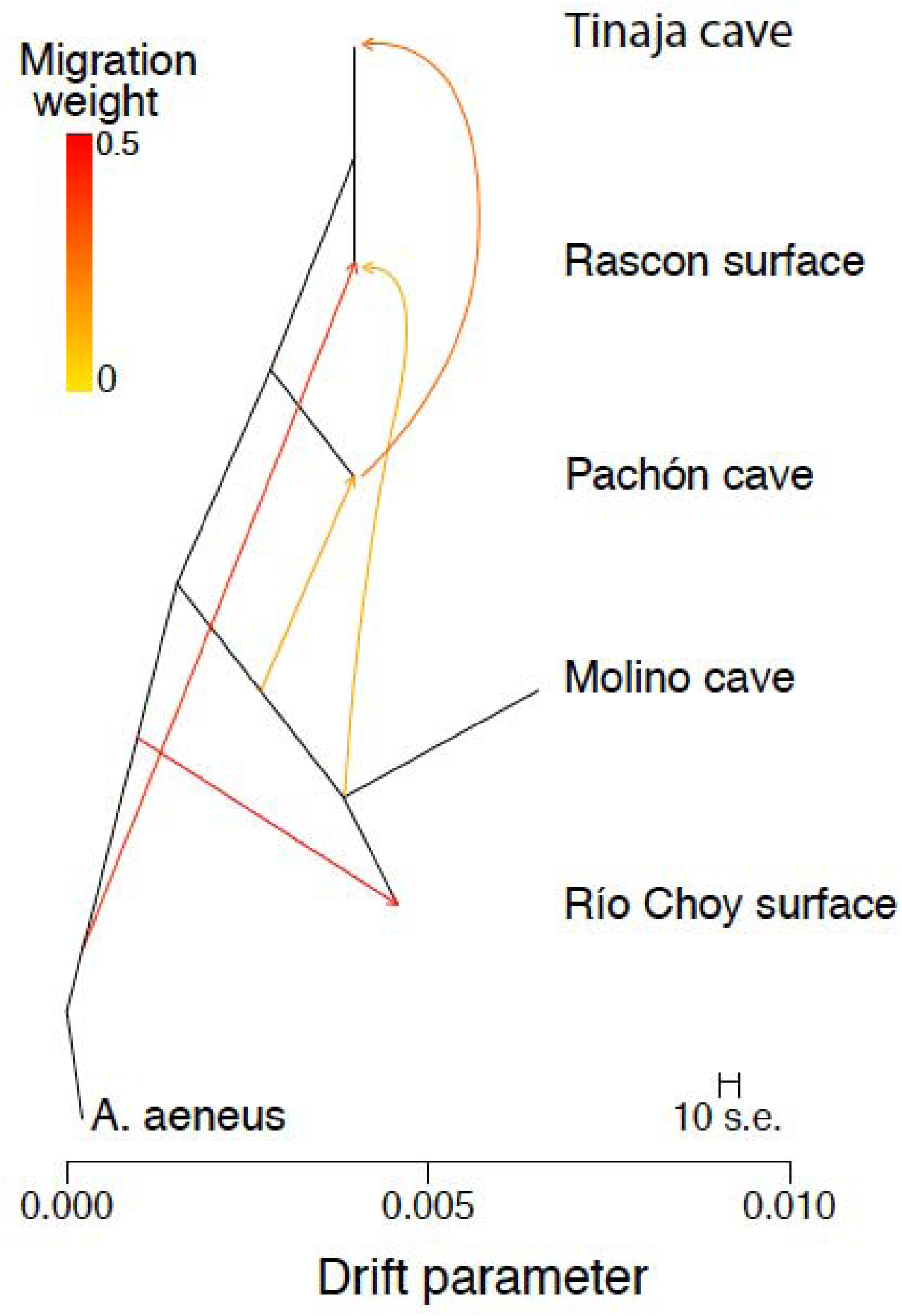
TreeMix population graph with five migration events. We see here patterns evident throughout our study, namely admixture between a lineage related to *A. aeneus* and Río Choy and Rascón, as well as exchange from the ancestor of Río Choy and Molino into Pachón and Rascón. Admixture from Pachón into Tinaja is also apparent. Analyses were conducted on the Biallelic SNP VCF file thinned to account for linkage between SNPs. This analysis excluded the four recently admixed individuals.

#### TreeMix

TreeMix allows us to visualize population relationships as a graph depicting directional admixture given a number of migration events, and uses the covariance structure of allele frequencies among populations rather than allele frequency difference correlations (e.g., *F*_*3*_ and *F*_*4*_). By examining multiple runs with different levels of migration, the TreeMix run with five migration events best explains the sample covariance (Figure S4).

TreeMix illustrates gene flow from the ancestral new lineage into both Rascón surface and Pachón cave (Figure 3; also suggested by *F*_*3*_ and *F*_*4*_ statistics), while providing evidence for gene flow from the lineage leading to *A. aeneus* samples to the Río Choy and Rascón lineages (also suggested by *F*_*3*_ and *F*_*4*_ statistics). Migration from Pachón cave into Tinaja cave is indicated (also supported by ∂a∂i modeling). However, the topology recovered by TreeMix places Pachón as the outgroup to (Rascón, Tinaja), thus, Pachón to Tinaja gene flow likely (partially) reflects this incorrect tree structure.

Care should be taken in interpreting the migration arrows drawn on the TreeMix result. First, we limited the number of migration edges to five, so further (possibly real) migration events are not represented. Second, any particular inferred admixture event is necessarily unidirectional and the source population is designated as the population with a migration weight ≤ 50%. Thus, while directionality is hard to pin down with these analyses, we can confidently report admixture among and between lineages and habitat types.

### Divergence times are younger than expected without accounting for migration

The levels of diversity and divergence allow us to calculate a simple estimate of population split times. This approach results in remarkably recent divergence time estimates. For example, we estimate the oldest split between old and new lineages of approximately 110,000 generations (Table 2; using π_4fold degen *A. aeneus*_ as a proxy for ancestral diversity, and representing the old-new lineage split by *d*_4fold degen Choy,Rascón_, (3.74 - 2.97) × 10^−3^ / (7 × 10^−9^)). Assuming the generation time of one year, the oldest estimate for the old and new split is more than an order of magnitude less than that estimated from the *cytochrome b* which places the divergence time between lineages between 5.7-7.5 mya (Ornelas-García *et al.* 2008). Similarly, we estimate that the split between new lineage cave (Molino) and new lineage surface (Río Choy) populations was approximately 50,000 generations ago ((3.32 - 3.00) × 10^−3^ / (7 × 10^−9^)) similar to a recent analyses (Fumey *et al.* 2018). However, other estimates using this method were unstable, suggesting that *A. aeneus* is not a good proxy for ancestral diversity in the old lineage and/or introgression obscures any realistic estimate of divergence time. Our results that suggest rampant gene flow (even with the exclusion of recently admixed individuals) indicate that true divergence times exceed those estimated from comparisons of *d*_XY_ and π. We, therefore, pursue a model-based estimate of population divergence time while accounting for introgression.

### Demographic modeling indicates cave populations are younger than expected when accounting for migration

Demographic modeling using ∂a∂i revealed extensive interdependence and contact for all populations studied. In all cases, models with multiple migration rates fit the population comparisons best, suggesting that some genomic regions are more recalcitrant to gene flow than others. For most pairwise population comparisons, the best-fitting models supported a period of isolation followed by a period of secondary contact (SC2m; Table S6). The only exceptions were the comparisons between Molino-Rascón and Pachón-Rascón that supported divergence with isolation (IM2m) slightly more than SC2m (Table S6). The distributions of divergence times for IM2m were multimodal (Molino-Rascón) or nearly flat (Pachón-Rascón) (Figure S9); thus, we present results from SC2m models that exhibit much tighter distributions for divergence estimates (Table 4; Figure S9). Results from all models for all replicates are provided (Table S8).

**Table 4.** ∂a∂i demographic modeling of divergence times between pairwise populations for 50 replicates per model. For population pairs that favored the IM2m model (e.g., isolation with migration and heterogeneity of migration across the genome), we also report results of SC2m (e.g., a period of isolation followed by secondary contact and heterogeneity of migration across the genome, black) which was the favored model for most pairwise population comparisons. Ts is the number of generations from population split to the start of secondary contact. Tsc is the number of generations from the start of secondary contact to the present. Ts+Tsc is the total divergence time in generations. Generation time is assumed to be six months to one year.

| Pop1 | Pop2 | Model | MedianTs | MinTs | MaxTs | MedianTsc | MinTsc | MaxTsc | MedianTS+TSC |
| --- | --- | --- | --- | --- | --- | --- | --- | --- | --- |
| Choy | Molino | SC2M | 91,270 | 0 | 1,031,510 | 71,268 | 0 | 2,061,383 | 162,537 |
| Choy | Pachón | SC2M | 174,483 | 0 | 5,894,387 | 6,690 | 0 | 1,944,141 | 181,173 |
| Choy | Rascón | SC2M | 241,747 | 7,018 | 263,584 | 14,928 | 0 | 274,878 | 256,675 |
| Choy | Tinaja | SC2M | 182,107 | 0 | 497,367 | 24,859 | 0 | 519,170 | 206,966 |
| Molino | Pachón | SC2M | 195,724 | 60 | 641,127 | 30,431 | 0 | 603,939 | 226,155 |
| Molino | Rascón | IM2M | 1,242,575 | 228,667 | 1,799,441 | NA | NA | NA | NA |
| Molino | Rascón | SC2M | 210,396 | 0 | 1,593,512 | 27,023 | 0 | 1,044,418 | 237,418 |
| Molino | Tinaja | SC2M | 119,177 | 0 | 563,918 | 34,819 | 0 | 290,502 | 153,996 |
| Pachón | Rascón | IM2M | 1,239,971 | 140,387 | 11,358,780 | NA | NA | NA | NA |
| Pachón | Rascón | SC2M | 148,702 | 27,575 | 655,675 | 12,560 | 0 | 644,435 | 161,262 |
| Pachón | Tinaja | SC2M | 102,245 | 0 | 261,637 | 13,277 | 4,358 | 137,507 | 115,522 |
| Rascón | Tinaja | SC2M | 170,279 | 0 | 405,939 | 20,370 | 0 | 399,350 | 190,650 |

Effective population sizes for the surface populations were an order of magnitude higher than for the cave populations, and Molino exhibited the lowest effective population size of all five populations (Table S9). Notably, even for cave populations, estimates of effective population size were an order of magnitude larger than previous estimates (Avise & Selander 1972; Bradic *et al.* 2012) and on par with mark-recapture census estimates (∼8500 fish in Pachon cave with wide 95% confidence interval of 1,279-18,283; Mitchell *et al.* 1977).

∂a∂i demographic modeling estimated deeper split times for populations than the divergence estimates calculated as T = (*d*_XY_ - π_ancestral_)/2μ (above) because they account for introgression that can artificially deflate these divergence estimates. The oldest divergence time between old and new lineages was estimated at 256,675 generations before present (Río Choy-Rascón), (Table 4, S10; Figure S9; using SC2m models for all comparisons). Importantly, both the model-independent method and the demographic estimates presented here reveal relatively similar divergence times between the old and new lineages (∼110k versus ∼257k generations ago), and these estimates differ substantially from the several million year divergence time obtained through mtDNA (Ornelas-García *et al.* 2008), unless we have dramatically underestimated generation time in the wild.

Cave-surface splits for both the new and old lineage comparisons were remarkably similar (Pachón - Rascón: 161,262 generations; Tinaja - Rascón: 190,650 generations; Molino - Río Choy: 162,537 generations). While the two old lineage cave populations (Pachón - Tinaja) are estimated to have split slightly more recently (115,522 generations before present), suggesting that colonization of one of the two old lineage caves was potentially from subterranean gene flow (as suggested by Espinasa & Espinasa 2015). Distributions and estimates across 50 replicates are given in Figure S9, Table S8, and Table S10.

Subterranean gene flow may be higher than surface to cave or between-lineage surface gene flow. The highest rates of gene flow are estimated between caves, and surprisingly, Molino cavefish exhibited a migration rate into Tinaja cave that is higher than Tinaja gene flow into Pachón cave. Surface fish gene flow rates into cave populations are also among the highest rates (especially Río Choy into all sampled caves). Notably, several caves also have relatively high rates of gene flow into surface populations (Molino into Río Choy and Tinaja into Rascón).

Heterogeneity in gene flow across the genome is also different across population pairs. Generally, more of the genome of cave-surface pairs (48%) seems to follow the lower migration rate than in cave-cave pairs (40%) and surface-surface pair (18%), suggesting the possibility that strong selection for habitat-specific phenotypes slows gene flow across the genome.

### Modeling of selection needed for cave alleles to reach high frequency

Similarly to that estimated by (Cartwright *et al.* 2017) selection coefficients of ∼0.01 needed are for cave-phenotype alleles to reach high frequencies in either Molino or Tinaja cave populations with a 12-locus additive model (Figure 4). Animations of allele frequencies at different levels of selection across the historical demography are provided in Supplementary materials. There is an increase in noise at the secondary contact period because the influx of new alleles. This has little effect overall because the migration rate is of the same magnitude as the mutation rate and may be because prior to secondary contact, the loci are typically fixed for either the cave or surface allele and after contact, the loci may become more polymorphic. The average selection coefficients appear to approximately match the pre-secondary contact average.

**Figure 4.**
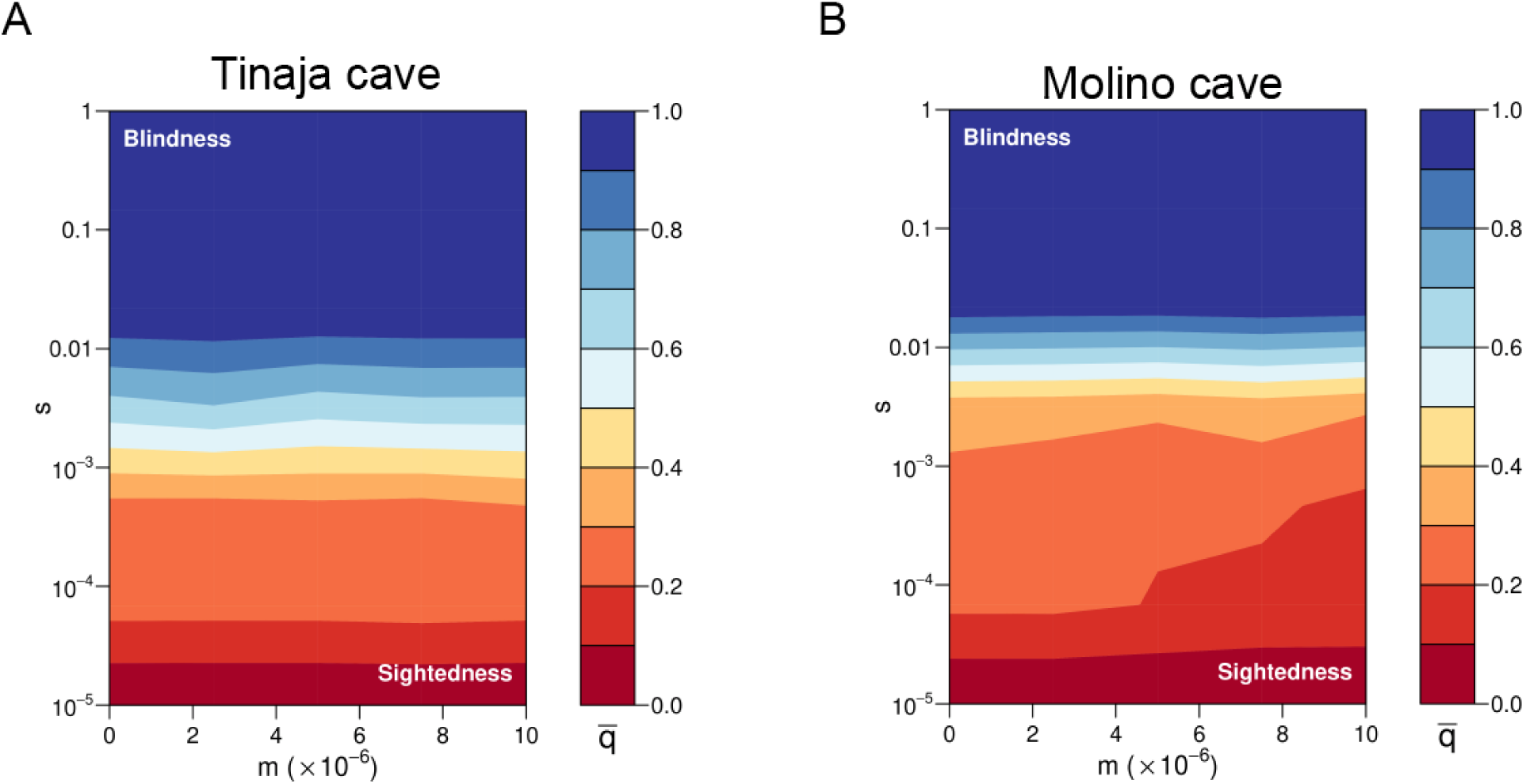
Selection coefficients required for cave alleles to reach high frequency (darker blue) in a 12-locus additive model. A. Tinaja. B. Molino. Shown is the final iteration of a model that takes into account isolation between cave and surface populations followed by secondary contact inline with demographic models presented in the paper. m = migration rate, s = selection coefficient.

In sum, our data support that cavefish population sizes are sufficiently large for selection to play a role shaping traits, gene flow is sufficiently common to impact repeated evolution, and cave-surface divergence is more recent than expected from mitochondrial gene trees.

### Candidate gene regions introgressed between caves

In light of the suspected gene flow among populations, we identified genomic regions that showed molecular evolution signatures consistent with gene flow between caves and genomic regions that are likely affected by cave-surface gene flow. There were 997 5kb regions (out of 208,354 total) that exhibited the lowest 5% of *d*_XY_ windows of the genome for each of the cave-cave comparisons (Pachón-Molino, Pachón-Tinaja, Molino-Tinaja). These windows were spread across 471 scaffolds and overlapped 500 genes. We highlight some of these 500 genes in the context of Ensembl phenotype data in Supplementary text (see also Table S11).

Several of these regions cluster with known QTLs (O’Quin & McGaugh 2015). We examined co-occurring QTLs for many traits on Linkage Group 2 and Linkage Group 17 (LG2, LG17, Table S12), which were two main regions highlighted in recent work on cave phenotypes (Yoshizawa *et al.* 2012b). We found that the co-occurring QTL on Linkage Group 2 harbored about 1.5-fold (odds ratio = 1.54, 95% CI =1.139-2.089) the number of regions with low-divergence among all three cave populations (44 windows with all three caves in lowest 5% of *d*_XY_ /6070 total windows that had data across these same scaffolds) relative to the total across the entire genome (997/208354). This suggests that this region with many co-occurring QTL was potentially transferred among cave. Notably, we did not find a similar pattern for linkage group 17 (proportion of windows in 5% lowest *d*_XY_ = 0.0048 for LG17 and 0.0048 across the entire genome). The region on Linkage Group 2 is not simply an area of high sequence conservation (which could account for low divergence among all three cave populations) because the mean divergence between surface-surface comparisons for this region is very similar to the genome-wide average (genome wide *d*_XY_ =0.0048, LG2 QTL region *d*_XY_ = 0.0048).

We find genes that look to be affected by gene flow between caves. We found 1004 genes that exhibited *d*_XY_ = 0 and another 370 genes with *d*_XY_ in the lowest 5% across the genome for the three pairwise comparisons of the caves, many with substantial divergence to at least one surface population (Table S13). We associated these genes with previously known QTL with linkage groups that follow (O’Quin *et al.* 2013) and a QTL database is provided (Table S7). One example is *fam136a* (*family with sequence similarity 136, member A*), which is under a QTL for body condition on Linkage Group 10. This gene is expressed in the hair cells of the crista ampullaris, an organ important for detecting rotation and acceleration, in the semicircular canals of the inner ear in the rat (Requena *et al.* 2014). All caves are identical except for a SNP that may have come from the Rascon population in one of the admixed Tinaja individuals. *fam136a* is in the top 10% most divergent genes by *d*_XY_ for Rascón surface – Pachón cave and Rascón surface – Tinaja cave populations (Molino-Rascón surface comparisons fall in the top 13% most divergent genes via *d*_XY_). Notably, Rascon and *A. aeneus* exhibit an E (glutamic acid) at 15aa (suggesting this is the ancestral state), whereas Río Choy and all caves exhibit a G (glycine). This amino acid switch is between a polar and a hydrophobic, thus not very functionally conserved. Such pattern could be produced by gene flow among caves or by gene flow of Río Choy with the cave populations.

## DISCUSSION

The role of admixture between diverging populations is increasingly apparent as the application of genomic analyses has become standard, and such gene flow shapes the study of repeated evolution (Colosimo *et al.* 2005; Cresko *et al.* 2004; Roesti *et al.* 2014; Van Belleghem *et al.* 2018; Welch & Jiggins 2014). Further, gene flow may create a signature that resembles parallel genetic divergence as effective migration is reduced in the same genomic regions for multiple ecotypic pairs upon secondary contact (Bierne *et al.* 2013; Rougemont *et al.* 2017). Thus, to understand the repeated origins of traits, an accurate understanding of the demography is required (Dasmahapatra *et al.* 2012; Rosenblum *et al.* 2014).

*Astyanax mexicanus* have been widely studied as a model of repeated evolution of wide-ranging traits (Elmer & Meyer 2011; Krishnan & Rohner 2017). Our data provide insight into some of the most pressing demographic questions needed to understand repeated evolution: 1) whether there are independent origins of *Astyanax* cavefish, 2) the age of cave invasions, 3) the amount of gene flow between populations, and 4) the strength of selection needed to shape traits. Genomic resequencing allowed an understanding not afforded by mitochondrial or reduced representation nuclear sequencing, and we were able to identify genes and a region of the genome potentially affected by gene flow between populations. Together, these findings provide a framework for understanding the repeated evolution of many complex, cave-derived traits.

### Cave populations are younger than previously estimated

Our estimates fit in well with previous suggestions that the caves were colonized in the late Pleistocene (Avise & Selander 1972; Fumey *et al.* 2018; Porter *et al.* 2007; Strecker *et al.* 2004) with both our *d*_XY_ – based divergence estimation and demographic modelling. As our work documents extensive gene flow, we favor the divergence time estimates provided by demographic models that incorporated estimates of migration. Demographic models estimate split time between the two lineages at approximately 257k generations ago, and estimates of cave-surface population splits are ∼161k - 191k generations ago. Notably, *Astyanax* is a species of fish with a limited distribution in northern latitudes, with its current most northern locality in the Edwards Plateau in Texas (Page & Burr 2011). These split times are consistent with cooler temperatures in Northern Mexico associated with glaciation playing some role in the colonization of the caves by *Astyanax mexicanus*, potentially as thermally-stable refugia (Cussac *et al.* 2009). Our nuclear genomic data suggest that the “old” and “new” lineage split is more recent than the main volcanic activity of the Trans-Mexican Volcanic Belt (3-12 Mya) which was thought to have separated these two lineages, as well as other lineages in other vertebrates (Ornelas-García *et al.* 2008). This geographic barrier likely has been breached by *Astyanax* multiple times and likely led to cave invasions with each migration (Gross 2012; Hausdorf *et al.* 2011; Strecker *et al.* 2012).

### Historic and contemporary gene flow between most populations is common

Several cave populations exhibit intermediate phenotypes and experience flooding during the rainy season (Espinasa, unpublished, Strecker *et al.* 2012), suggesting that intermediate phenotypes are the result of admixture between cavefish and surface fish swept into caves during flooding (Avise & Selander 1972; Bradic *et al.* 2012). However, it has been suggested that surface fish and hybrids are too maladapted to survive and spawn in caves (Coghill *et al.* 2014; Hausdorf *et al.* 2011; Strecker *et al.* 2012), and that cavefish populations with intermediate troglomorphic phenotypes represent more recent cave invasions, rather than hybrids (Hausdorf *et al.* 2011; Strecker *et al.* 2012). Despite evidence from microsatellites (Bradic *et al.* 2012; Panaram & Borowsky 2005), mitochondrial capture (Dowling *et al.* 2002; Ornelas-García & Pedraza-Lara 2015; Yoshizawa *et al.* 2012a), and haplotype sharing of candidate loci (but see Espinasa *et al.* 2014c; Gross & Wilkens 2013), there was still uncertainty in the literature regarding the frequency of gene flow between *Astyanax* cavefish and surface fish and the role gene flow may play in adaptation to the cave environment (Coghill *et al.* 2014; Espinasa & Borowsky 2001; Hausdorf *et al.* 2011; Strecker *et al.* 2012).

Our work demonstrates recent and historical gene flow between cave and surface populations both within and between lineages (Figures 2, 3). This result is also suggested by past work which showed most genetic variance is within individuals (Table 2 in (Bradic *et al.* 2012)) and very few private alleles (i.e., alleles specific to a population) were present among cave populations (Figure 3b in (Bradic *et al.* 2012)). Notably, many of the methods we employed take into account incomplete lineage sorting, are robust to non-equilibrium demographic scenarios, and detect recent admixture (< 500 generations ago) (Patterson *et al.* 2012). In all our analyses the hybridization detected may not be between these sampled populations directly, but unsampled populations/lineages related to them (Patterson *et al.* 2012).

One of the most intensely studied cave populations is Pachón. Our data of Pachón cavefish hybridization with the new lineage is expected from past field observations and molecular data. Only extremely troglomorphic fish were found early (1940’s-1970’s; Avise & Selander 1972; Mitchell *et al.* 1977) and late (1996-2000, (Dowling *et al.* 2002)) surveys, though, phenotypically intermediate fish were observed in 1986-1988, as well as in 2008 (Borowsky, unpublished). Thus, subterranean introgression from a nearby cave population may cause transient complementation of phenotypes or hybridization with surface fish (from flooding or human-introduction) may contribute to the presence of intermediate-phenotype fish in the cave (Langecker *et al.* 1991; Wilkens & Strecker 2017). Indeed past studies suggested gene flow between Pachón and the new lineage populations, as Pachón mtDNA clusters with new lineage populations (Dowling *et al.* 2002; Ornelas-García *et al.* 2008; Ornelas-García & Pedraza-Lara 2015; Strecker *et al.* 2003; Strecker *et al.* 2012). Interestingly and divergent from past studies, Pachón mitochondria group with old-lineage populations in the most recent analysis (Coghill *et al.* 2014). Continued intense scrutiny of other caves may reveal similar fluctuations in phenotypes and genotypes.

The most surprising signal of gene flow in our data is seen in the ∂a∂i analyses, in which the signal of Molino-Tinaja exchange is similar to that from Pachón-Tinaja, suggesting subterranean gene flow between all three caves, despite substantial geographic distances (the entrances are separated by >100km; Mitchell *et al.* 1977). While our genome-wide ancestry proportion data suggest some exchange of Molino with Tinaja (Figure S5-S7), we have no other evidence to corroborate this signal. Thus, the results from population demography could be picking up a signal of Molino with Tinaja that is actually driven by Pachón - new lineage hybridization and subsequent Tinaja – Pachón hybridization or Tinaja - Río Choy hybridization.

Notably, many species of troglobites other than *Astyanax* inhabit caves spanning from southern El Abra to Sierra de Guatemala, and some were able to migrate among these areas within the last 12,000 years (Espinasa *et al.* 2014a). Due to the extensive connectivity and the span of geological changes throughout the hydrogeological history, it is likely that there have been ample opportunities for troglobites, including *Astyanax*, to migrate across much of the El Abra region (Espinasa & Espinasa 2015).

Across many systems genomic data has revealed reticulate evolution is much more common than previously thought (Abbott *et al.* 2016; Arnold & Kunte 2017; Brandvain *et al.* 2014; Dasmahapatra *et al.* 2012; Geneva *et al.* 2015; Malinsky *et al.* 2017; McGaugh & Noor 2012; Rougemont *et al.* 2017), and the ability of populations to maintain phenotypic differences despite secondary contact and hybridization is increasingly appreciated (Arnold & Kunte 2017; Fitzpatrick *et al.* 2015; Malinsky *et al.* 2017; Payseur & Rieseberg 2016). Thus, despite gene flow from the surface into caves, it is entirely possible that cave-like phenotypes can be maintained. Indeed, gene flow may help sustain cave populations from the effects of inbreeding (Åkesson *et al.* 2016; Ellstrand & Rieseberg 2016; Fitzpatrick *et al.* 2016; Frankham 2015; Kronenberger *et al.* 2017; Whiteley *et al.* 2015) or catalyze adaptation to the cave environment (sensu Clarkson *et al.* 2014; Meier *et al.* 2017; Richards & Martin 2017). Our data indicate extensive hybridization between sampled lineages and suggest that cavefish are poised to be a strong contributor to understanding the role gene flow may play in repeated evolutionary adaptation.

### Candidate genes shaped by gene flow

In some cases, we have evidence that particular alleles were transferred between caves, but are highly diverged from surface populations (Table S11, S13). This suggests that gene flow between caves may hasten adaptation to the cave environment and suggests that repeated evolution in this system may, in part, rely on standing genetic variation. Notably, scaffolds under the co-localizing QTL on Linkage Group 2 for many traits exhibit 1.5 fold enrichment for 5kb regions with lowest divergence across caves relative to genome-wide, suggesting parts of this QTL have spread throughout caves.

For genes with high similarity among caves and substantial divergence between cave and surface, many appeared to be involved in classic cavefish traits such as pigmentation and eye development/morphology (*mdkb*, *atp6v0ca*, *ube2d2l*, *usp3*, *cln8*) and circadian functioning (*usp2a*) (Table S13). Additional common annotations suggest future areas of trait investigation, namely cardiac-related phenotypes (*mmd2*, *cyp26a1*, *tbx3a*, *tnfsf10*, *alx1*, *ptgr1*) as well as inner ear phenotypes (*ncs1a, dlx6a, fam136a*) (Table S13). Importantly, the sensory hair cells of the inner ear are homologous with sensory hair cells of the neuromasts of the lateral line (Fay & Popper 2000), therefore, these genes may be impacting mechanoreception or sound reception.

### Effective population sizes are much larger than expected and weak selection can drive cave phenotypes

Previous estimates of very small effective population sizes in cave populations of *A. mexicanus* suggested drift and relaxed selection shaped cave-derived phenotypes (e.g., Lahti *et al.* 2009). The estimates of nucleotide diversity and *N*_*e*_ provided here (Tables 1, 2, S9) indicated that positive selection coefficients need not be extreme to drive cave-derived traits (Akashi *et al.* 2012; Charlesworth 2009). To put these values in perspective, the cave populations have an average genetic diversity (which is proportional to *N*_*e*_) similar to humans and the surface populations exhibit a similar genetic diversity to zebrafish (Leffler et al. 2012).

Our observations support theoretical and empirical results that selection likely shaped cavefish phenotypes (Borowsky 2015; Cartwright *et al.* 2017; Moran *et al.* 2015). Simulations using demographic parameters from Molino and Tinaja cave populations suggest that selection coefficients across 12 additive loci (patterned after the number of eye-related QTL (O’Quin & McGaugh 2015)) need to be above 0.01 to bring cave-alleles to very high frequencies (Figure 4). Such selection coefficients are often found driving selective sweeps in natural systems (Nair *et al.* 2003; Rieseberg & Burke 2001; Schlenke & Begun 2004; Wootton *et al.* 2002).

### Diverging lineages should be examined with reticulation methods and absolute divergence metrics

Our genome-wide data allowed a unique perspective that was not available in past studies. First, our data represent an empirical demonstration of how well-supported phylogenetic reconstructions can be misleading. When using genome-scale data with maximum likelihood, bootstrap values are not a measure of the number of sites that support a particular phylogeny, though, this is often how they are interpreted (Yang & Rannala 2012). Rather, if a genome-scale dataset is slightly more supportive of a particular topology, maximum likelihood will find that topology consistently and exhibit high levels of bootstrap support (Yang & Rannala 2012). With the cavefish, a previous RADseq study suggested relatively strongly supported branches (Coghill *et al.* 2014), however, a much more complex evolutionary history with substantial reticulation among lineages was revealed here. Examination of recently diverged taxa with reticulation methods (e.g., Phylonet, *F*_*4*_, *F*_*3*_, TreeMix) can ensure that phylogenetic reconstructions do not provide unwarranted confidence in a bifurcating tree.

Second, our dataset is yet another empirical demonstration of poor performance of pairwise F_ST_ in estimating population relationships when diversity is highly heterogeneous among populations (Figure 5, Tables 2, S14; Fumey *et al.* 2018; Jakobsson *et al.* 2013). When diversity is highly heterogeneous, this can give the false impression that low-diversity populations are highly divergent, when in reality the lower diversity drives greater pairwise F_ST,_ not higher absolute divergence (Charlesworth 1998). These limitations of F_ST_ are especially important to appreciate in systems like the cavefish where diversity is highly heterogeneous across populations. Further, it is often suggested that high F_ST_ translates to low gene flow, but violations of the assumptions are common (Cruickshank & Hahn 2014). In the case of Molino (a very low diversity population), high F_ST_ values were taken to indicate relative isolation from other populations with little gene flow (Bradic *et al.* 2012), and we show here that is not the case. Indeed, pairwise F_ST_ among surface fish populations is the lowest and pairwise F_ST_ among caves is the highest, yet, absolute divergence between surface populations is slightly higher than absolute divergence among caves (Figure 5, Tables 2, S14). Future molecular ecology work should assay diversity and interpret pairwise F_ST_ accordingly or use absolute measures of divergence in conjunction with pairwise F_ST_ (Charlesworth 1998; Cruickshank & Hahn 2014; Noor & Bennett 2009; Ritz & Noor 2016).

**Figure 5.**
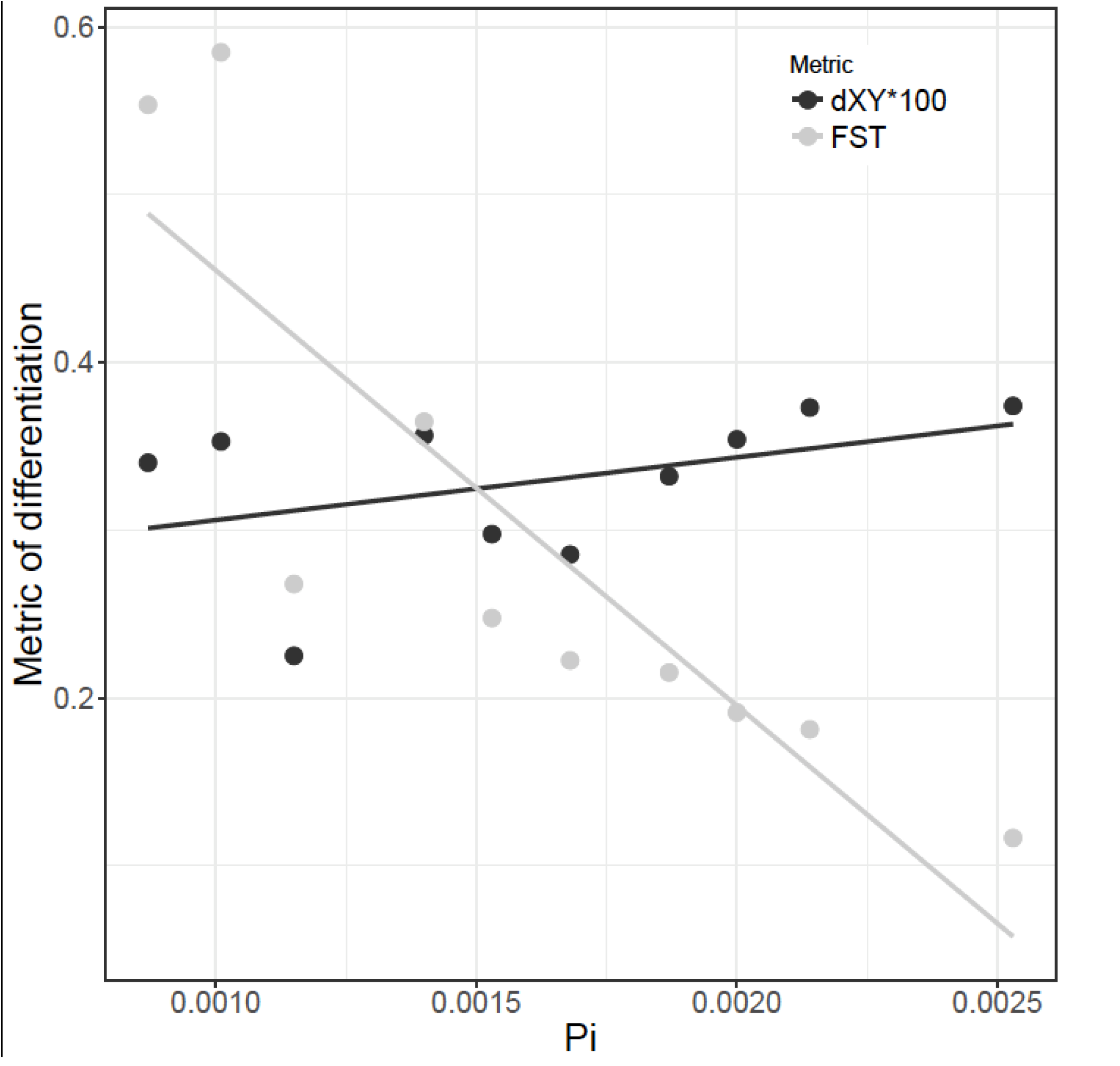
Pairwise population comparisons using F_ST_ and average π _pop1-pop2_ at fourfold degenerate sites (filled circles) or *d*_XY_ (open circles) averaged across the genome using non-admixed individuals. There is no correlation between F_ST_ and *d*_XY_, but a strong correlation between F_ST_ and average π _pop1-pop2_. Thus, the heterogeneity in π is driving F_ST_, not absolute divergence.

### Conclusion

In conclusion, our results suggest investigations of repeated evolution of cave-derived traits should take into account hybridization between lineages, between cave and surface fish, and between caves. Due to gene flow, best practices to assign the putative ancestral character state of cave-derived traits include comparing cavefish phenotypes to multiple surface populations and a distantly related outgroup with no evidence of admixture (e.g. *Astyanax bimaculatus*, Ornelas-García *et al.* 2008). Complementation crosses and molecular studies remain essential for understanding evolutionary origins for each cave-derived trait (Borowsky 2008b; Gross *et al.* 2009; O’Quin *et al.* 2015; Protas *et al.* 2006; Wilkens 1971; Wilkens & Strecker 2003).

We expect future work with greatly expanded sampling (Beerli 2004; Hellenthal *et al.* 2014; Pease & Hahn 2015; Slatkin 2005) and ecological parameterization of the caves to provide a more comprehensive view of the ultimate drivers of demography of *Astyanax mexicanus* cavefish. Interestingly, many caves are nutrient-limited which is suggested to be one of the largest impediments to surface fish survival in the cave environment (Espinasa *et al.* 2014b; Moran *et al.* 2014). One of the emerging hypotheses is that high food availability allows surface-like fish to persist longer in the cave environment and enhances the probability for cave-surface hybridization (Mitchell *et al.* 1977; Strecker *et al.* 2012). Future studies evaluating hybridization levels in relation to food-availability or food-predictability are an important next step to examine important drivers of cave phenotypes. We look forward to these future studies, and their potential to elucidate how demography and environmental factors impact repeated adaptive evolution (sensu Rosenblum *et al.* 2014).

## Acknowledgments

Fish were collected under CONAPESCA permit PPF/DGOPA - 106 / 2013 to Claudia Patricia Ornelas García and SEMARNAT permit 02241 to Ernesto Maldonado. We thank the Mexican government for providing the collecting permit to R.B. in 2008 (DGOPA.00570.288108-0291). For 2002, the collection permit to R.B. was from fisheries department #01.01.02.613.03.1799 Molino samples were obtained under Mexican permit 040396-213-05. Animal care protocol numbers include #05-1235 by the New York University Animal Welfare Committee (UAWC) to R.B., UMD R-17-77 to WRJ, and UNAM animal care protocol to POG NOM-062-ZOO-1999. This work was supported by NIH grant 2R24OD011198-04A1 to WCW, The Genome Institute at Washington University School of Medicine, Cave Research Foundation Graduate Student Research Grant to BMC, a grant from the Eppley Foundation for Research to SEM, 5R01EY014619-08 to WRJ, and 1R01GM127872-01 to SEM and ACK. Raw sequence data were submitted to the SRA. Project Accession Number: SRP046999, Bioproject: PRJNA260715. We appreciate the resources provided by the Minnesota Supercomputing Institute, without which this work would not be possible.

---

**Table S1.**
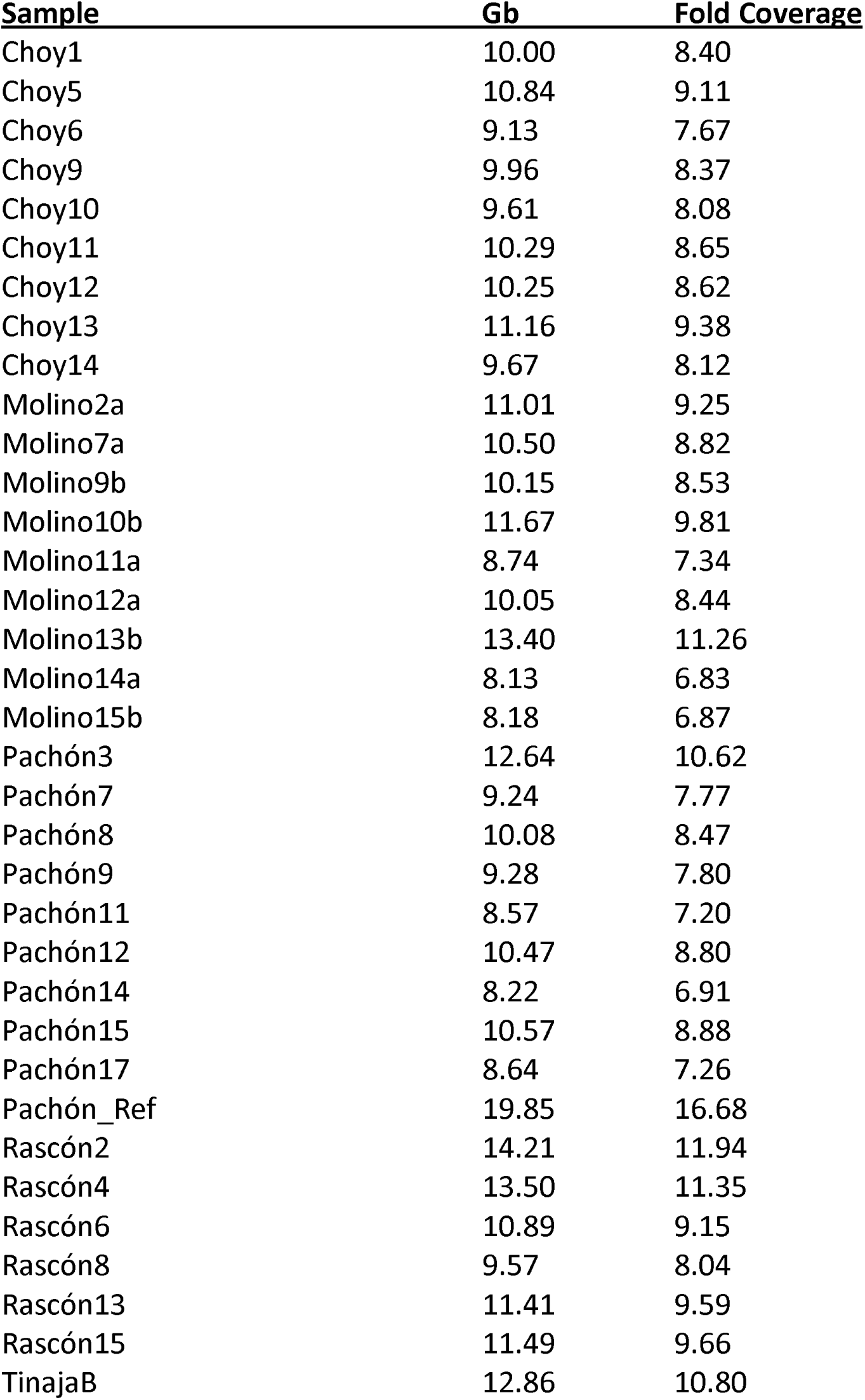
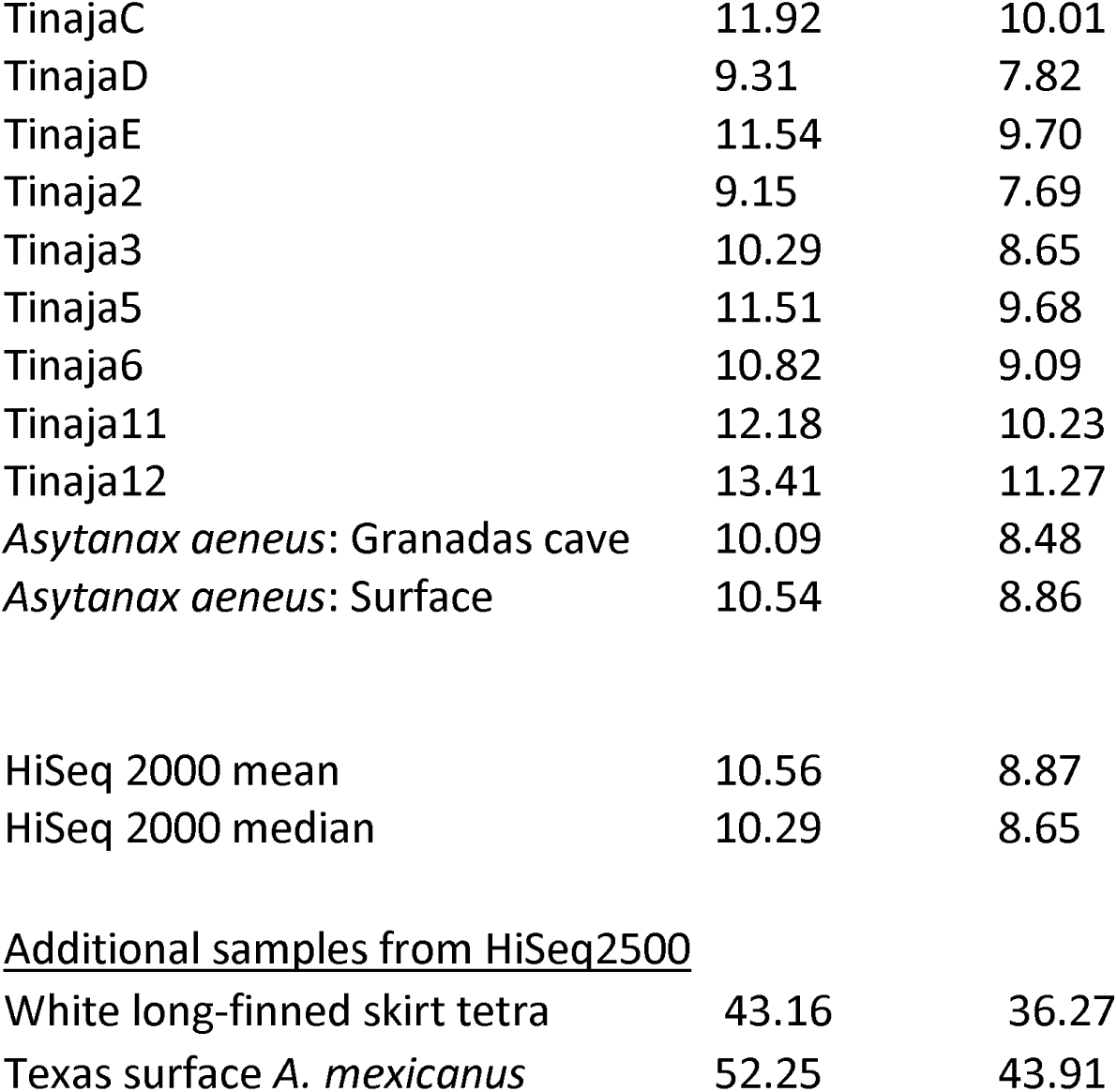
Sample name and sequence coverage for the core samples analyzed here after filtering for quality and removing adaptors. Approximate fold coverage is given using the genome size estimate of 1.19 Gb (McGaugh *et al.* 2014). All samples were sequenced with Illumina DNA nano library preps and v3 chemistry using the HiSeq 2000. Pachón_Ref were single-end reads from the genome project. Several individuals were sequenced on HiSeq 2500 rather than HiSeq 2000. These include the outgroup white long-finned skirt tetra (*Gymnocorymbus ternetzi*) and Texas surface *A. mexicanus*.

| **Sample** | **Gb** | **Fold Coverage** |
| --- | --- | --- |
| Choy1 | 10.00 | 8.40 |
| Choy5 | 10.84 | 9.11 |
| Choy6 | 9.13 | 7.67 |
| Choy9 | 9.96 | 8.37 |
| Choy10 | 9.61 | 8.08 |
| Choy11 | 10.29 | 8.65 |
| Choy12 | 10.25 | 8.62 |
| Choy13 | 11.16 | 9.38 |
| Choy14 | 9.67 | 8.12 |
| Molino2a | 11.01 | 9.25 |
| Molino7a | 10.50 | 8.82 |
| Molino9b | 10.15 | 8.53 |
| Molino10b | 11.67 | 9.81 |
| Molino11a | 8.74 | 7.34 |
| Molino12a | 10.05 | 8.44 |
| Molino13b | 13.40 | 11.26 |
| Molino14a | 8.13 | 6.83 |
| Molino15b | 8.18 | 6.87 |
| Pachón3 | 12.64 | 10.62 |
| Pachón7 | 9.24 | 7.77 |
| Pachón8 | 10.08 | 8.47 |
| Pachón9 | 9.28 | 7.80 |
| Pachón11 | 8.57 | 7.20 |
| Pachón12 | 10.47 | 8.80 |
| Pachón14 | 8.22 | 6.91 |
| Pachón15 | 10.57 | 8.88 |
| Pachón17 | 8.64 | 7.26 |
| Pachón_Ref | 19.85 | 16.68 |
| Rascón2 | 14.21 | 11.94 |
| Rascón4 | 13.50 | 11.35 |
| Rascón6 | 10.89 | 9.15 |
| Rascón8 | 9.57 | 8.04 |
| Rascón13 | 11.41 | 9.59 |
| Rascón15 | 11.49 | 9.66 |
| TinajaB | 12.86 | 10.80 |
| TinajaC | 11.92 | 10.01 |
| TinajaD | 9.31 | 7.82 |
| TinajaE | 11.54 | 9.70 |
| Tinaja2 | 9.15 | 7.69 |
| Tinaja3 | 10.29 | 8.65 |
| Tinaja5 | 11.51 | 9.68 |
| Tinaja6 | 10.82 | 9.09 |
| Tinaja11 | 12.18 | 10.23 |
| Tinaja12 | 13.41 | 11.27 |
| *Asytanax aeneus*: Granadas cave | 10.09 | 8.48 |
| *Asytanax aeneus*: Surface | 10.54 | 8.86 |
| HiSeq 2000 mean | 10.56 | 8.87 |
| HiSeq 2000 median | 10.29 | 8.65 |
| Additional samples from HiSeq2500 |  |  |
| White long-finned skirt tetra | 43.16 | 36.27 |
| Texas surface *A. mexicanus* | 52.25 | 43.91 |

**Table S2.** Sample name and alignment coverage before and after removing PCR duplicates, along with the proportion of reads removed as duplicates. Pachón_Ref were single-end reads from the genome project. Several individuals were sequenced on HiSeq 2500 rather than HiSeq 2000. These include the outgroup white long-finned skirt tetra (*Gymnocorymbus ternetzi*) and Texas surface *A. mexicanus*.

| **Sample** | **Merged** | **Duplicate Removed** | **Proportion Duplicate** |
| --- | --- | --- | --- |
| **Reads** |  |  |  |
| Choy1 | 9.19 | 8.80 | 0.042 |
| Choy5 | 9.95 | 9.56 | 0.039 |
| Choy6 | 8.38 | 8.01 | 0.044 |
| Choy9 | 9.14 | 8.76 | 0.042 |
| Choy10 | 8.82 | 8.49 | 0.037 |
| Choy11 | 9.41 | 9.00 | 0.044 |
| Choy12 | 9.42 | 9.03 | 0.041 |
| Choy13 | 10.26 | 9.80 | 0.045 |
| Choy14 | 8.88 | 8.51 | 0.042 |
| Molino2a | 10.13 | 9.50 | 0.062 |
| Molino7a | 9.67 | 9.17 | 0.052 |
| Molino9b | 9.35 | 8.86 | 0.052 |
| Molino10b | 10.74 | 10.13 | 0.057 |
| Molino11a | 8.04 | 7.59 | 0.056 |
| Molino12a | 9.24 | 8.70 | 0.058 |
| Molino13b | 12.32 | 11.51 | 0.066 |
| Molino14a | 7.47 | 7.06 | 0.055 |
| Molino15b | 7.52 | 7.12 | 0.053 |
| Pach3 | 11.78 | 11.28 | 0.042 |
| Pach7 | 8.60 | 8.25 | 0.041 |
| Pach8 | 9.37 | 8.98 | 0.042 |
| Pach9 | 8.64 | 8.28 | 0.042 |
| Pach11 | 7.98 | 7.66 | 0.040 |
| Pach12 | 9.76 | 9.32 | 0.045 |
| Pach14 | 7.65 | 7.34 | 0.041 |
| Pach15 | 9.86 | 9.46 | 0.041 |
| Pach17 | 8.04 | 7.71 | 0.041 |
| Pach_Ref | 17.74 | 13.36 | 0.247 |
| Rascón2 | 13.15 | 12.59 | 0.043 |
| Rascón4 | 12.47 | 11.92 | 0.044 |
| Rascón6 | 10.06 | 9.61 | 0.045 |
| Rascón8 | 8.63 | 8.28 | 0.041 |
| Rascón13 | 10.51 | 10.05 | 0.044 |
| Rascón15 | 10.60 | 10.11 | 0.046 |
| TinajaB | 11.84 | 11.22 | 0.052 |
| TinajaC | 10.96 | 10.39 | 0.052 |
| TinajaD | 8.56 | 8.15 | 0.048 |
| TinajaE | 10.63 | 10.10 | 0.050 |
| Tinaja2 | 8.42 | 8.01 | 0.049 |
| Tinaja3 | 9.41 | 8.94 | 0.050 |
| Tinaja5 | 7.27 | 6.93 | 0.047 |
| Tinaja6 | 9.88 | 9.38 | 0.051 |
| Tinaja11 | 11.21 | 10.64 | 0.051 |
| Tinaja12 | 12.35 | 11.70 | 0.053 |
| *Asytanax aeneus*: Surface | 9.61 | 9.06 | 0.057 |
| *Asytanax aeneus*: Granadas cave | 9.23 | 8.78 | 0.049 |
| HiSeq 2000 mean | 9.83 | 9.28 | 0.052 |
| HiSeq 2000 median | 9.42 | 9.02 | 0.046 |
| Additional samples from HiSeq2500 |  |  |  |
| White long-finned skirt tetra | 18.94 | 15.03 | 0.206 |
| Texas surface *A. mexicanus* | 47.68 | 42.85 | 0.101 |

**Table S3.**
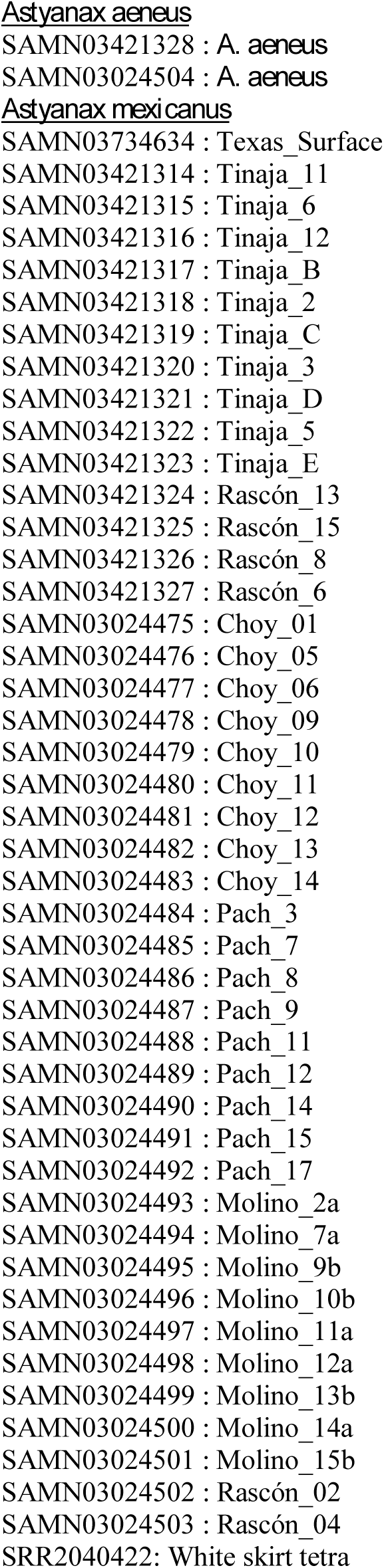
Sequence Read Archive accession numbers for raw data reads are located under Bioproject: PRJNA260715.

**Table S4.** Parameters used to filter variant call set: sites for which these parameters were true were marked and removed from downstream analyses.

| **Parameter Name** | **Filter Values**<br>**for SNPs** | **Filter Values**<br>**for Mixed/Indels** |
| --- | --- | --- |
| QualByDepth (QD) | < 2.0 | < 2.0 |
| FisherStrand (FS) | > 60.0 | > 200.0 |
| RMSMappingQuality (MQ) | < 40.0 | -- |
| MappingQualityRankSumTest (MQRankSum) | < -12.5 | -- |
| ReadPosRankSumTest (ReadPosRankSum) | < -8.0 | < -20.0 |

**Table S5.** Description of the parameters used in the ∂a∂i model. All parameter estimates are reported with respect to an internally-defined effective population size, Nref (multiply by the value in the Scaling Factor column to convert to real-world values). Not all parameters are used by all models.

| Parameter | Description | Scaling Factor | Starting Value |
| --- | --- | --- | --- |
| **N1** | Ne of population 1 | $N_{\text{ref}}$ | 1 |
| **N2** | Ne of population 2 | $N_{\text{ref}}$ | 1 |
| **Ts** | Time from split to present (SI, IM models) or secondary contact (SC models) | $2N_{\text{ref}}$ | 1 |
| **Tsc** | Time from secondary contact to present | $2N_{\text{ref}}$ | 0.1 |
| **Tam** | Time from end of ancestral migration to split | $2N_{\text{ref}}$ | 0.1 |
| **m21** | Migration from population 1 to population 2 | $1/(2N_{\text{ref}})$ | 5 |
| **m12** | Migration from population 2 to population 1 | $1/(2N_{\text{ref}})$ | 5 |
| **mi21** | Migration from population 1 to population 2 in “islands” | $1/(2N_{\text{ref}})$ | 0.5 |
| **mi12** | Migration from population 2 to population 1 in “islands” | $1/(2N_{\text{ref}})$ | 0.5 |
| **p** | Proportion of genome in “islands” | None | 0.5 |

**Table S6:** Model selection for ∂a∂i demographic modeling using weighted AIC values across the 50 replicates for each model. Higher values indicate the favored model. The seven models are as follows: SI - strict isolation, SC - secondary contact, IM - isolation with migration, AM - ancestral migration, SC2M - secondary contact with two migration rates, IM2M - isolation with two migration rates, and AM2M - ancestral migration with two migration rates (see Figure S9 of (Tine *et al.* 2014)).

| **Pop1** | **Pop2** | **BestModel** | **SI** | **SC** | **IM** | **AM** | **SC2M** | **IM2M** | **AM2M** |
| --- | --- | --- | --- | --- | --- | --- | --- | --- | --- |
| Choy | Molino | SC2M | 0 | 0 | 0 | 0 | 0.46 | 0.41 | 0.13 |
| Choy | Pachon | SC2M | 0 | 0.47 | 0 | 0 | 0.53 | 0 | 0 |
| Choy | Rascón | SC2M | 0 | 0 | 0 | 0 | 0.42 | 0.31 | 0.27 |
| Choy | Tinaja | SC2M | 0 | 0.02 | 0 | 0 | 0.58 | 0.34 | 0.60 |
| Molino | Pachon | SC2M | 0 | 0 | 0 | 0 | 0.51 | 0.41 | 0.08 |
| Molino | Rascón | IM2M | 0 | 0 | 0 | 0 | 0.24 | 0.63 | 0.12 |
| Molino | Tinaja | SC2M | 0 | 0 | 0 | 0 | 0.48 | 0.32 | 0.20 |
| Pachon | Rascón | IM2M | 0 | 0 | 0 | 0 | 0.34 | 0.60 | 0.06 |
| Pachon | Tinaja | SC2M | 0 | 0 | 0 | 0 | 0.78 | 0.15 | 0.06 |
| Rascón | Tinaja | SC2M | 0 | 0 | 0 | 0 | 0.51 | 0.37 | 0.12 |

**Table S7:** Median and ranges of effective population size estimates for each population calculated from all SC2m models and replicates which involved that population.

| **Population** | **Median Ne** | **Min Ne** | **Max Ne** |
| --- | --- | --- | --- |
| **Río Choy** | 258,643.48 | 219,187.97 | 706,222.19 |
| **Molino** | 7,335.23 | 1,803.14 | 18,661.97 |
| **Pachón** | 31,755.67 | 3,052.04 | 46,023.74 |
| **Rascón** | 173,816.85 | 128,892.74 | 597,166.07 |
| **Tinaja** | 30,522.15 | 11,144.70 | 42,453.10 |

**Table S8:**
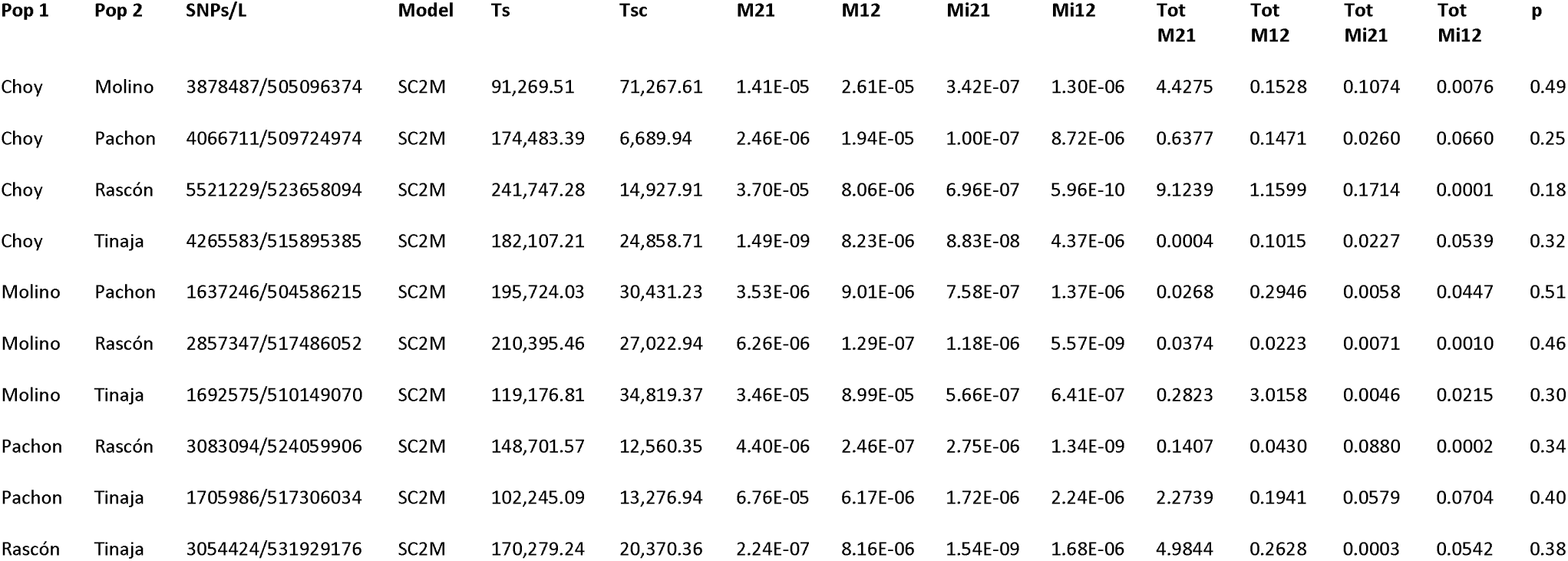
Median estimates of population divergence and migration rate parameters for best-fitting models across 50 replicates. All parameter estimates are scaled to real-world values. L is the number of sites, including invariant sites, that were used to generate the 2D SFS for each comparison. Ts is the number of generations from population split to the start of secondary contact. Tsc is the number of generations from the start of secondary contact to the present. M21 and M12 are the proportions of migrants from population 1 and 2, respectively. Mi21 and Mi12 are migrant proportions, but for genomic regions that may be recalcitrant to migration. P is the proportion of the genome that follows Mi21 and Mi12 migration rates. Values for all models for all replicates are given as a separate supplementary document. Columns with migration rates starting with “Tot” are the median number of chromosomes that are migrant, per generation (i.e., the median migrant proportion multiplied by the population size).

| Pop 1 | Pop 2 | SNPs/L | Model | Ts | Tsc | M21 | M12 | Mi21 | Mi12 | Tot M21 | Tot M12 | Tot Mi21 | Tot Mi12 | p |
| --- | --- | --- | --- | --- | --- | --- | --- | --- | --- | --- | --- | --- | --- | --- |
| Choy | Molino | 3878487/505096374 | SC2M | 91,269.51 | 71,267.61 | 1.41E-05 | 2.61E-05 | 3.42E-07 | 1.30E-06 | 4.4275 | 0.1528 | 0.1074 | 0.0076 | 0.49 |
| Choy | Pachon | 4066711/509724974 | SC2M | 174,483.39 | 6,689.94 | 2.46E-06 | 1.94E-05 | 1.00E-07 | 8.72E-06 | 0.6377 | 0.1471 | 0.0260 | 0.0660 | 0.25 |
| Choy | Rascón | 5521229/523658094 | SC2M | 241,747.28 | 14,927.91 | 3.70E-05 | 8.06E-06 | 6.96E-07 | 5.96E-10 | 9.1239 | 1.1599 | 0.1714 | 0.0001 | 0.18 |
| Choy | Tinaja | 4265583/515895385 | SC2M | 182,107.21 | 24,858.71 | 1.49E-09 | 8.23E-06 | 8.83E-08 | 4.37E-06 | 0.0004 | 0.1015 | 0.0227 | 0.0539 | 0.32 |
| Molino | Pachon | 1637246/504586215 | SC2M | 195,724.03 | 30,431.23 | 3.53E-06 | 9.01E-06 | 7.58E-07 | 1.37E-06 | 0.0268 | 0.2946 | 0.0058 | 0.0447 | 0.51 |
| Molino | Rascón | 2857347/517486052 | SC2M | 210,395.46 | 27,022.94 | 6.26E-06 | 1.29E-07 | 1.18E-06 | 5.57E-09 | 0.0374 | 0.0223 | 0.0071 | 0.0010 | 0.46 |
| Molino | Tinaja | 1692575/510149070 | SC2M | 119,176.81 | 34,819.37 | 3.46E-05 | 8.99E-05 | 5.66E-07 | 6.41E-07 | 0.2823 | 3.0158 | 0.0046 | 0.0215 | 0.30 |
| Pachon | Rascón | 3083094/524059906 | SC2M | 148,701.57 | 12,560.35 | 4.40E-06 | 2.46E-07 | 2.75E-06 | 1.34E-09 | 0.1407 | 0.0430 | 0.0880 | 0.0002 | 0.34 |
| Pachon | Tinaja | 1705986/517306034 | SC2M | 102,245.09 | 13,276.94 | 6.76E-05 | 6.17E-06 | 1.72E-06 | 2.24E-06 | 2.2739 | 0.1941 | 0.0579 | 0.0704 | 0.40 |
| Rascón | Tinaja | 3054424/531929176 | SC2M | 170,279.24 | 20,370.36 | 2.24E-07 | 8.16E-06 | 1.54E-09 | 1.68E-06 | 4.9844 | 0.2628 | 0.0003 | 0.0542 | 0.38 |

**Table S9.** Pairwise population comparisons using F_ST_ (lower triangle) and average π _pop1-pop2_ at fourfold degenerate sites (upper triangle) averaged across the genome using non-admixed individuals. Comparisons to Molino are among the highest F_ST_ values, and we suspect this is due to the low diversity in Molino. Likewise, Choy and Rascón are the most divergent in *d*_XY_, but exhibit the highest π and lowest F_ST_.

|  | *A. aeneus* | Río Choy | Molino | Pachón | Rascón | Tinaja |
| --- | --- | --- | --- | --- | --- | --- |
| *A. aeneus* | -- | 0.00299 | 0.00186 | 0.00199 | 0.00252 | 0.00213 |
| Río Choy | 0.1680 | -- | 0.00187 | 0.00200 | 0.00253 | 0.00214 |
| Molino | 0.6507 | 0.2151 | -- | 0.00087 | 0.00140 | 0.00101 |
| Pachón | 0.5158 | 0.1913 | 0.5534 | -- | 0.00153 | 0.00115 |
| Rascón | 0.2396 | 0.1165 | 0.3646 | 0.2476 | -- | 0.00168 |
| Tinaja | 0.4968 | 0.1812 | 0.5847 | 0.2678 | 0.2223 | -- |

**Figure S1.**
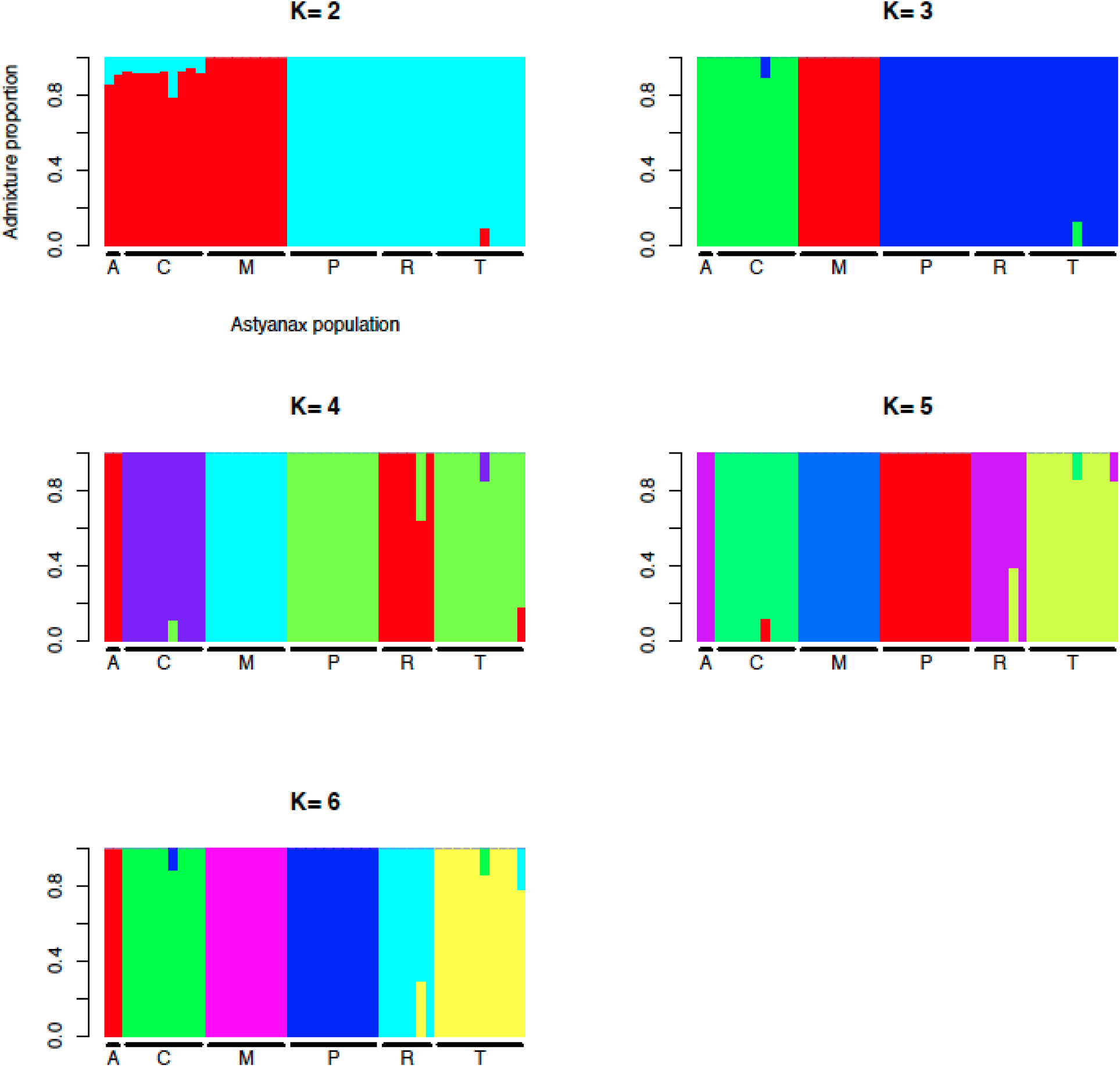
ADMIXTURE analysis suggests contemporary gene flow between “old” and “new” lineages and cave and surface populations using different cluster sizes. Analysis was performed on a thinned Biallelic SNP VCF. A = *Astyanax aeneus*, N = 2; C = Río Choy, N = 10; M = Molino cave, N = 9; P = Pachón cave, N = 10; R = Rascón surface, N = 6; T = Tinaja cave, N = 10.

**Figure S2.**
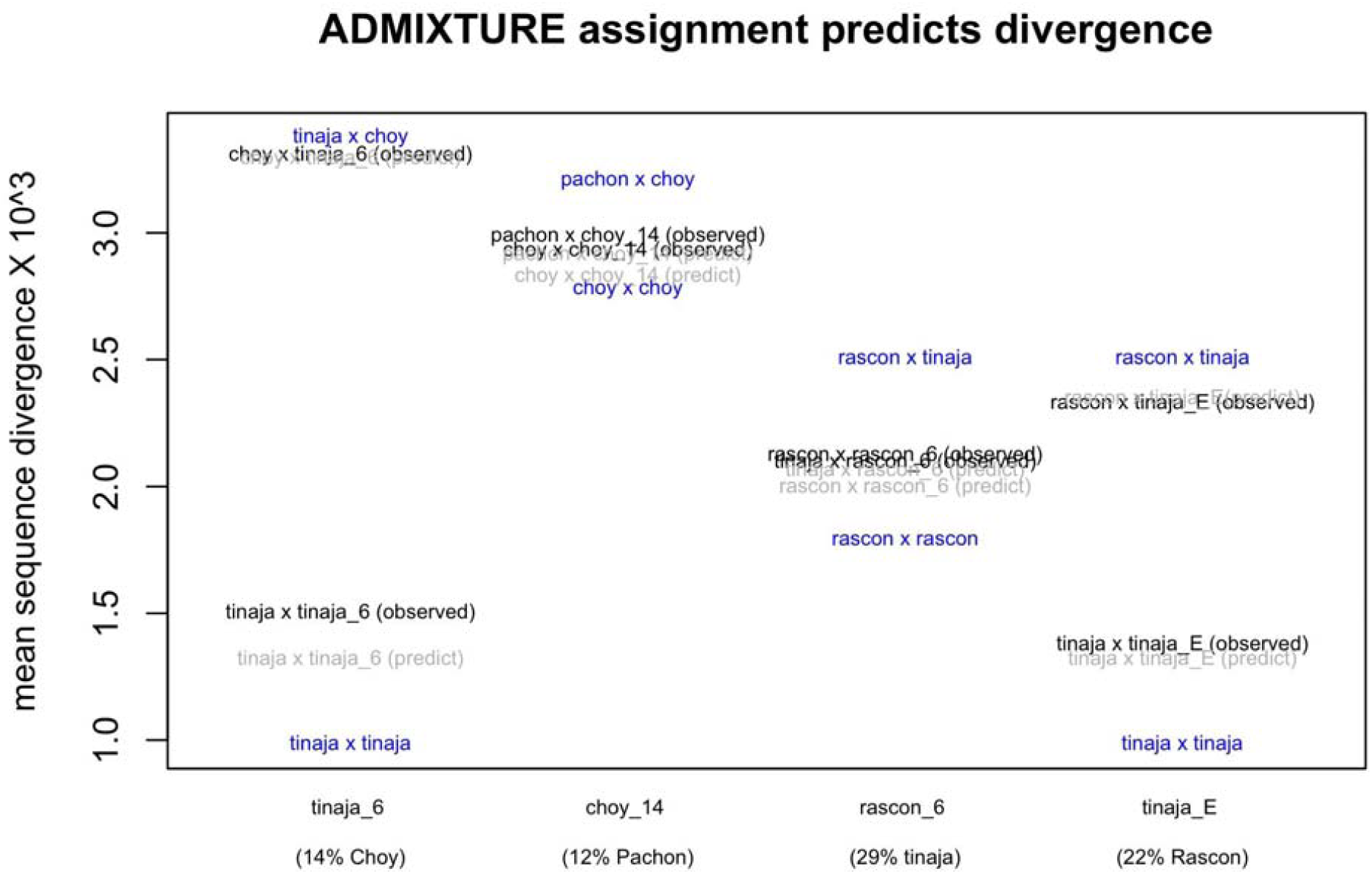
ADMIXTURE assignment predicts divergence. To validate our finding of early generation hybrids, we model divergence between a focal admixed sample its two parental species as diversity within and divergence between the parental species weighted by the admixture proportion inferred. For example, because ADMIXTURE predicts that sample Tinaja_E inherits 22% of its ancestry from Tinaja and 78% of its ancestry from Rascón, we predict that its divergence from Tinaja will be 0.78 π_Tinaja_ + 0.22 *d*_*Tinaja x Rascón*_, and that its divergence from Rascón will be 0.78 *d*_*Tinaja x Rascón*_ + 0.22 π_Rascón_. These predicted divergence values of 1.3 × 10^3^ and 2.3 × 10^3^ are quite close to the observed divergence values of 1.4 × 10^3^ and 2.3 × 10^3^, respectively. We visually present all such comparisons below.

**Figure S3.**
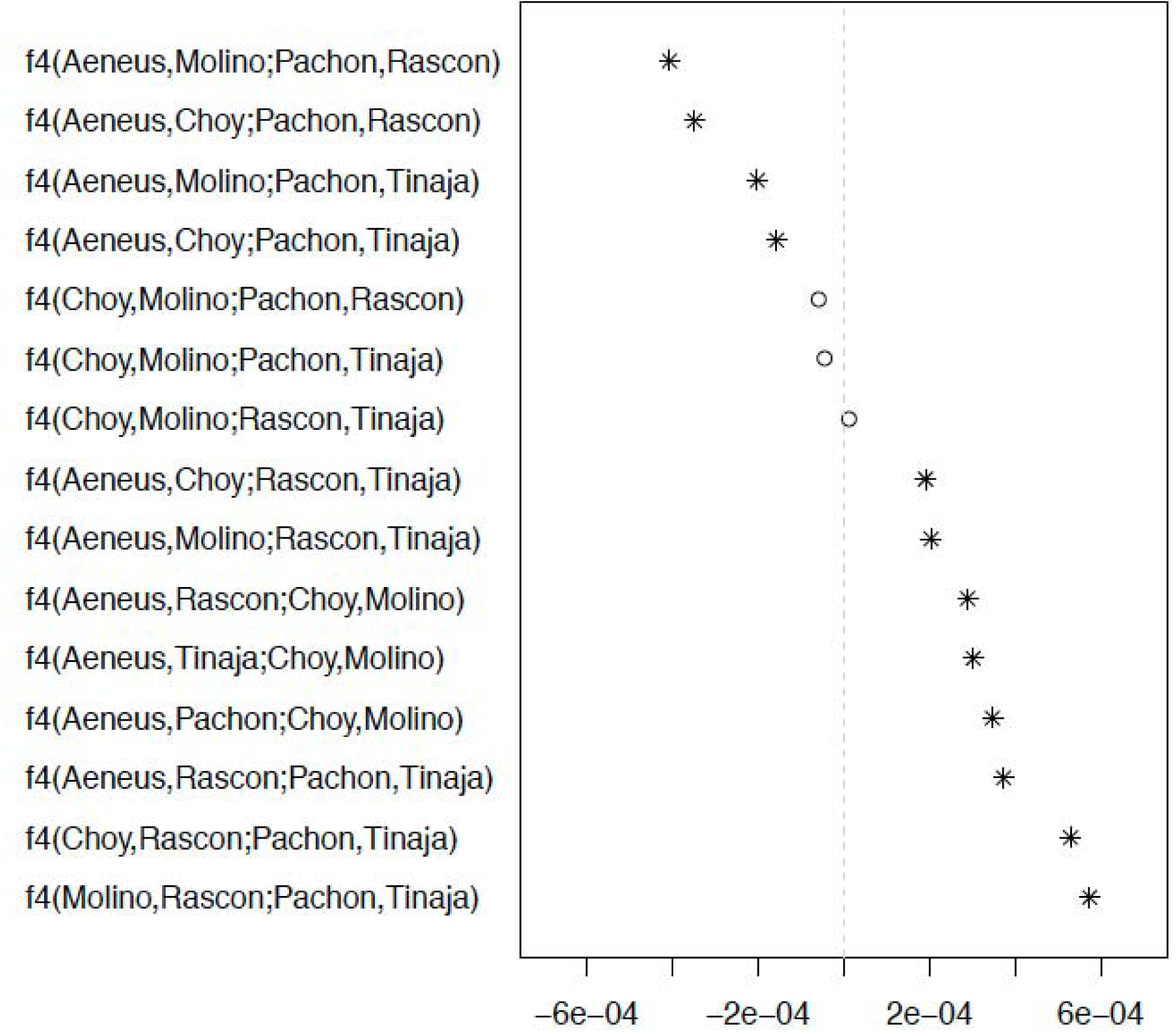
*F*_*4*_ statistics as calculated in TreeMix. Asterisks indicate a significant Z-score. For those newick trees where the asterisk falls in the negative zone, admixture between either the two outermost or the two inner-most is favored. For those newick trees where the asterisk falls in a positive zone, admixture between the first and third or second and fourth taxa is supported. Putatively recently admixed individuals were removed from the analysis.

**Figure S4.**
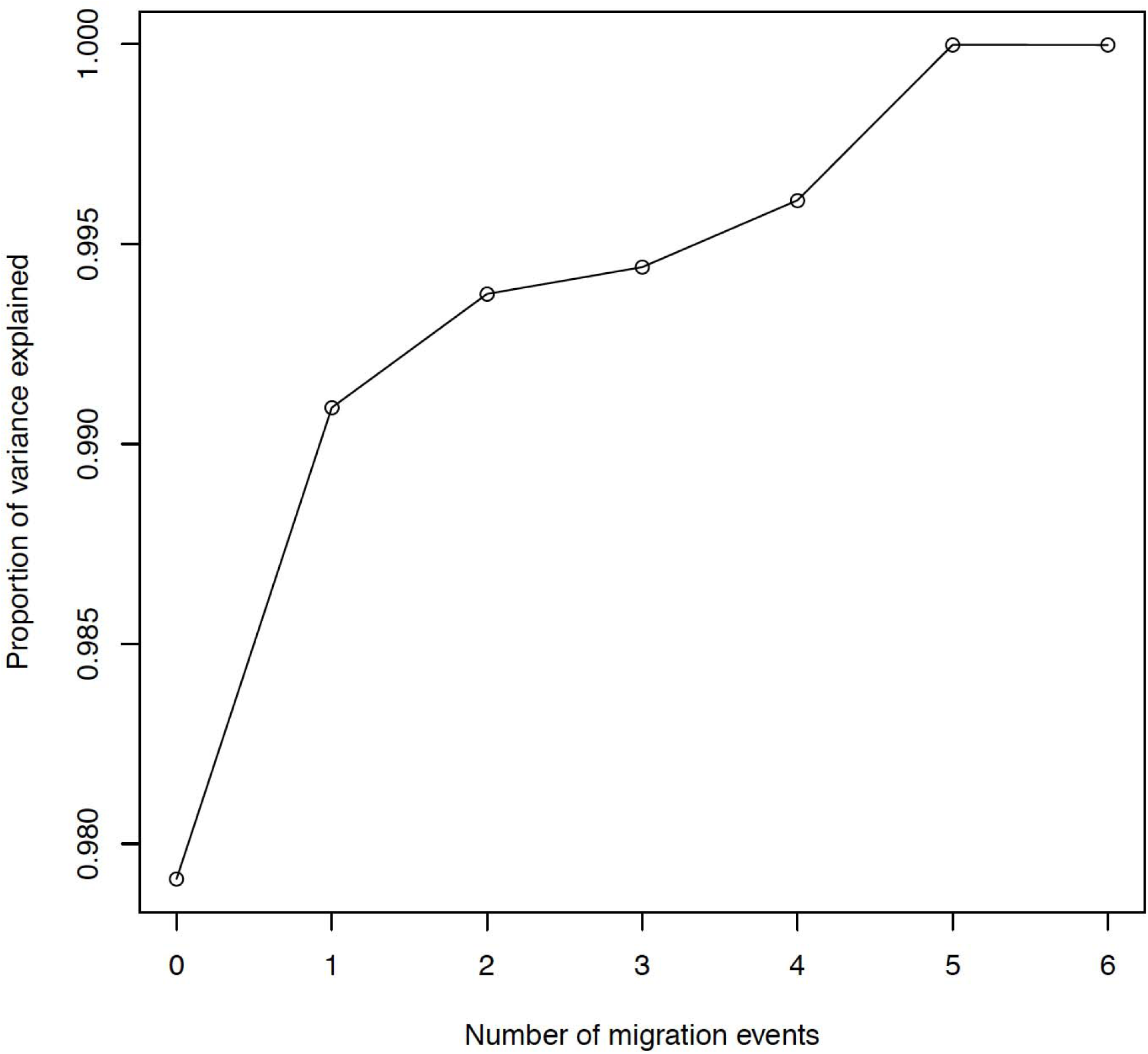
TreeMix proportion of variance explained by number of migration events. We calculated the proportion of variance explained as the observed sample covariance / expected sample covariance. There is clearly a plateau from five to six migration events, indicating that five events sufficiently explain the observed covariance matrix.

**Figure S5.**
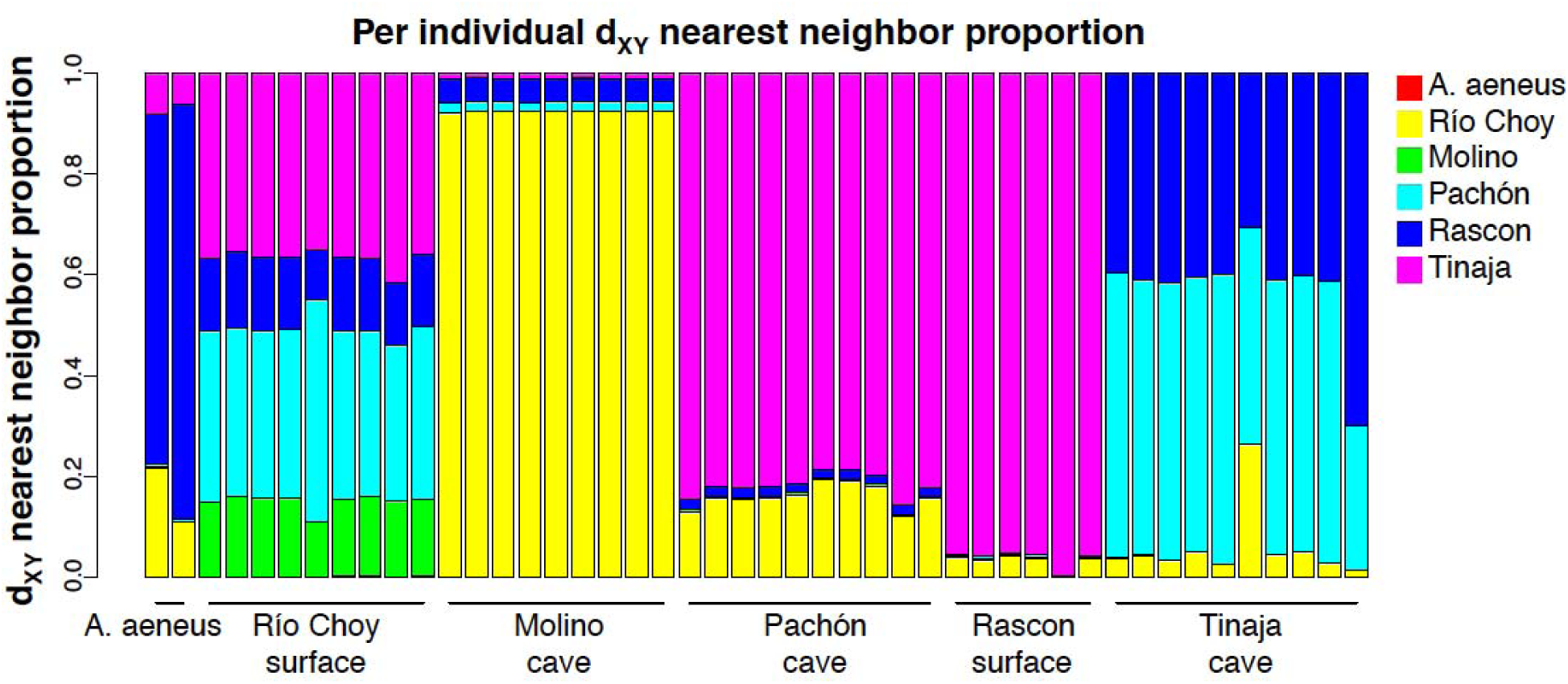
Per-individual estimated copying proportions from *d*_XY_-based chromosome painting for non-overlapping 20kb windows across the genome. We restrict these proportions to chunks that are at least 20kb and have unambiguous population affinity (i.e., for each window the assigned population is consistent and the only population with minimum *d*_XY_ to the focal individual).

**Figure S6.** Examination of length distribution of ancestry chunks per individual. 5 kb windows of average pairwise nucleotide differences (e.g., *d*_XY_) were assigned to their nearest-neighbor population in the sampled populations. **We include this as a separate PDF**.

**Figure S7.** Pairwise individual *d*_XY_ for each individual (on y-axis) for each 5kb window that show that population being the minimum *d*_XY_ distance from the focal population (listed as the title). For example, 5kb windows that show *A. aeneus* as the nearest *d*_XY_ neighbor to Choy, are much more divergent than when the other four populations are Choy’s nearest *d*_XY_ neighbor. Populations are abbreviated by their first letter. **We include this for every individual as a separate PDF.**

**Figure S8.** Best topology from using 1430 loci in PhyloNet v3.6.1. MCMC_SEQ incorporating reticulate evolution and incomplete lineage sorting. **We include this for every individual as a separate PDF. Analyses shown below were conducted with four recently admixed individuals removed and *apriori* information included.**

**Figure S9.**
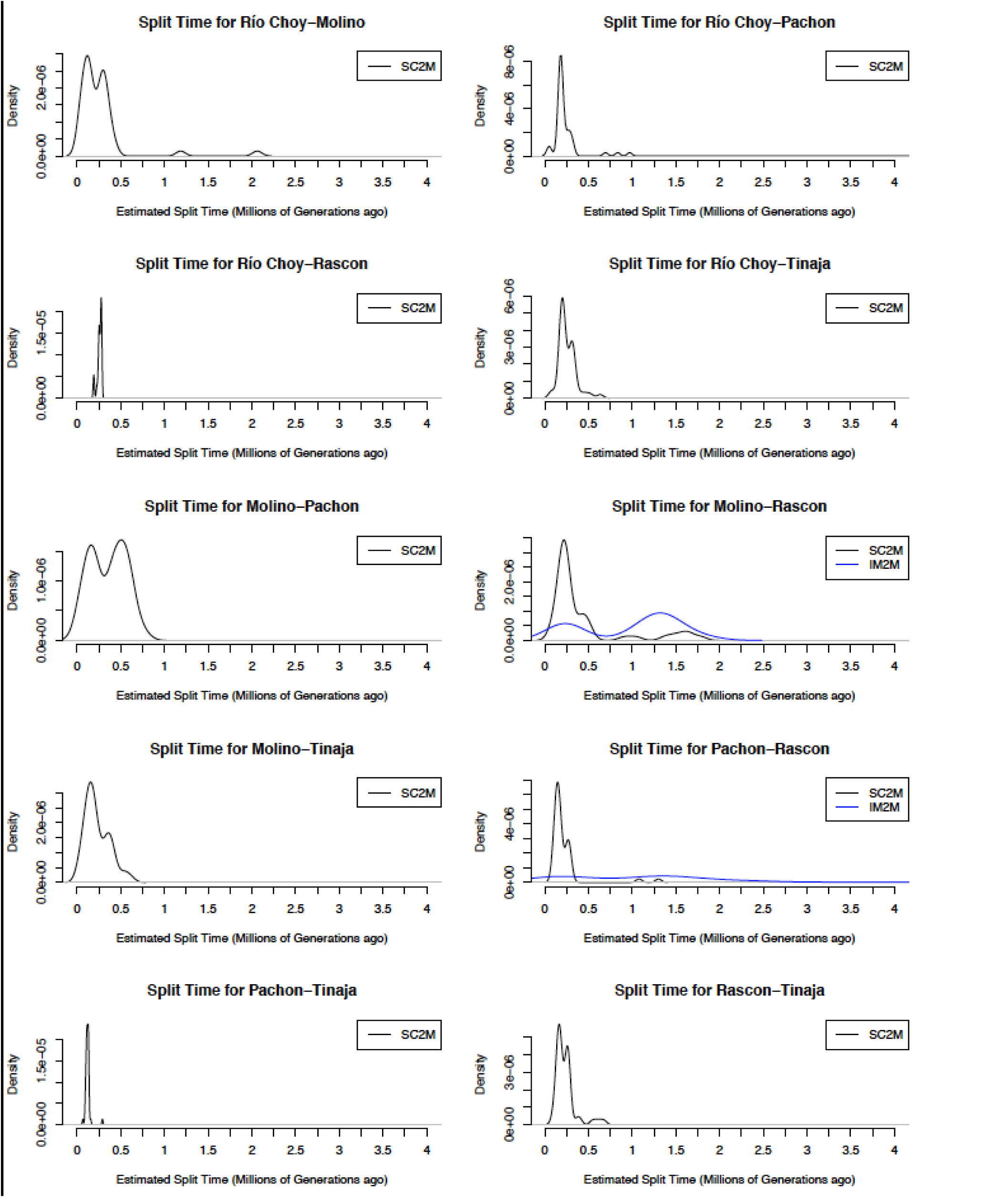
∂a∂i demographic modeling of divergence times between pairwise populations for 50 replicates per model. For population pairs that favored the IM2m model (e.g., isolation with migration and heterogeneity of migration across the genome, blue), we also plotted the results of SC2m (e.g., a period of isolation followed by secondary contact and heterogeneity of migration across the genome, black) which was the favored model for most pairwise population comparisons.

## References cited

Abbott RJ, Barton NH, Good JM (2016) Genomics of hybridization and its evolutionary consequences. Molecular ecology 25, 2325–2332.

Agrawal AA (2017) Toward a predictive framework for convergent evolution: integrating natural history, genetic mechanisms, and consequences for the diversity of life. The American Naturalist 190, S1–S12.

Åkesson M, Liberg O, Sand H, et al. (2016) Genetic rescue in a severely inbred wolf population. Molecular ecology 25, 4745–4756.

Alexander DH, Lange K (2011) Enhancements to the ADMIXTURE algorithm for individual ancestry estimation. BMC bioinformatics 12, 1.

Alexander DH, Novembre J, Lange K (2009) Fast model-based estimation of ancestry in unrelated individuals. Genome research 19, 1655–1664.

Arnold ML, Kunte K (2017) Adaptive genetic exchange: A tangled history of admixture and evolutionary innovation. Trends in Ecology & Evolution.

Aspiras AC, Rohner N, Martineau B, Borowsky RL, Tabin CJ (2015) Melanocortin 4 receptor mutations contribute to the adaptation of cavefish to nutrient-poor conditions. Proceedings of the National Academy of Sciences 112, 9668–9673.

Auwera GA, Carneiro MO, Hartl C, et al. (2013) From FastQ data to highlJconfidence variant calls: The genome analysis toolkit best practices pipeline. Current protocols in bioinformatics, 11.10. 11-11.10. 33.

Avise JC, Selander RK (1972) Evolutionary genetics of cave-dwelling fishes of the genus Astyanax. Evolution, 1–19.

Beerli P (2004) Effect of unsampled populations on the estimation of population sizes and migration rates between sampled populations. Molecular ecology 13, 827–836.

Bibliowicz J, Alié A, Espinasa L, et al. (2013) Differences in chemosensory response between eyed and eyeless Astyanax mexicanus of the Rio Subterráneo cave. EvoDevo 4, 25–25.

Bierne N, Gagnaire P-A, David P (2013) The geography of introgression in a patchy environment and the thorn in the side of ecological speciation. Current Zoology 59, 72–86.

Borowsky R (2008a) Astyanax mexicanus, the blind Mexican cave fish: A model for studies in development and morphology. Cold Spring Harbor Protocols 2008, pdb. emo107.

Borowsky R (2008b) Restoring sight in blind cavefish. Current Biology 18, R23–R24.

Borowsky R (2015) Regressive evolution: Testing hypotheses of selection and drift. In: Biology and Evolution of the Mexican Cavefish (ed. Keene A, Yoshizawa, M. McGaugh, S. E.), p. 93. Elsevier, San Diego.

Bradic M, Beerli P, León FG-d, Esquivel-Bobadilla S, Borowsky R (2012) Gene flow and population structure in the Mexican blind cavefish complex (Astyanax mexicanus). BMC evolutionary biology 12, 9.

Bradic M, Teotónio H, Borowsky RL (2013) The population genomics of repeated evolution in the blind cavefish Astyanax mexicanus. Molecular biology and evolution 30, 2383–2400.

Brandvain Y, Kenney AM, Flagel L, Coop G, Sweigart AL (2014) Speciation and introgression between Mimulus nasutus and Mimulus guttatus. PLoS genetics 10, e1004410.

Browning SR, Browning BL (2007) Rapid and accurate haplotype phasing and missing-data inference for whole-genome association studies by use of localized haplotype clustering. The American Journal of Human Genetics 81, 1084–1097.

Capella-Gutiérrez S, Silla-Martínez JM, Gabaldón T (2009) trimAl: a tool for automated alignment trimming in large-scale phylogenetic analyses. Bioinformatics 25, 1972–1973.

Charlesworth B (1998) Measures of divergence between populations and the effect of forces that reduce variability. Molecular biology and evolution 15, 538–543.

Clarkson CS, Weetman D, Essandoh J, et al. (2014) Adaptive introgression between Anopheles sibling species eliminates a major genomic island but not reproductive isolation. Nature communications 5.

Coghill LM, Hulsey CD, Chaves-Campos J, García de Leon FJ, Johnson SG (2014) Next generation phylogeography of cave and surface Astyanax mexicanus. Molecular Phylogenetics and Evolution 79, 368–374.

Colosimo PF, Hosemann KE, Balabhadra S, et al. (2005) Widespread parallel evolution in sticklebacks by repeated fixation of ectodysplasin alleles. Science 307, 1928–1933.

Cresko WA, Amores A, Wilson C, et al. (2004) Parallel genetic basis for repeated evolution of armor loss in Alaskan threespine stickleback populations. Proc Natl Acad Sci U S A 101, 6050–6055.

Cruickshank TE, Hahn MW (2014) Reanalysis suggests that genomic islands of speciation are due to reduced diversity, not reduced gene flow. Molecular ecology 23, 3133–3157.

Cussac V, Fernández DA, Gómez SE, López HL (2009) Fishes of southern South America: a story driven by temperature. Fish Physiology and Biochemistry 35, 29–42.

Dasmahapatra KK, Walters JR, Briscoe AD, et al. (2012) Butterfly genome reveals promiscuous exchange of mimicry adaptations among species. Nature 487, 94.

DePristo MA, Banks E, Poplin R, et al. (2011) A framework for variation discovery and genotyping using next-generation DNA sequencing data. Nature genetics 43, 491–498.

Dowling TE, Martasian DP, Jeffery WR (2002) Evidence for multiple genetic forms with similar eyeless phenotypes in the blind cavefish, Astyanax mexicanus. Molecular biology and evolution 19, 446–455.

Duboué ER, Keene AC, Borowsky RL (2011) Evolutionary convergence on sleep loss in cavefish populations. Current Biology 21, 671–676.

Ellegren H (2014) Genome sequencing and population genomics in non-model organisms. Trends in Ecology & Evolution 29, 51–63.

Ellstrand NC, Rieseberg LH (2016) When gene flow really matters: gene flow in applied evolutionary biology. Evolutionary applications 9, 833–836.

Elmer KR, Meyer A (2011) Adaptation in the age of ecological genomics: insights from parallelism and convergence. Trends in Ecology and Evolution 26, 298–306.

Espinasa L, Bartolo ND, Newkirk CE (2014a) DNA sequences of troglobitic nicoletiid insects support Sierra de El Abra and the Sierra de Guatemala as a single biogeographical area: Implications for Astyanax. Subterranean Biology 13, 35.

Espinasa L, Bibliowicz J, Jeffery WR, Rétaux S (2014b) Enhanced prey capture skills in Astyanax cavefish larvae are independent from eye loss. EvoDevo 5, 35.

Espinasa L, Borowsky RB (2001) Origins and relationship of cave populations of the blind Mexican tetra, Astyanax fasciatus, in the Sierra de El Abra. Environmental Biology of Fishes 62, 233–237.

Espinasa L, Centone DM, Gross JB (2014c) A contemporary analysis of a loss-of-function of the oculocutaneous albinism type II (Oca2) allele within the Micos Astyanax cave fish population. Speleobiology Notes 6, 48–54.

Espinasa L, Espinasa M (2015) Hydrogeology of caves in the Sierra de El Abra region. In: Biology and Evolution of the Mexican Cavefish. (eds. Keene A, Yoshizawa M, McGaugh S), pp. 41–58. Elsevier San Diego.

Espinasa L, Rivas-Manzano P, Pérez HE (2001) A new blind cave fish population of genus Astyanax: geography, morphology and behavior. In: The biology of hypogean fishes, pp. 339–344. Springer.

Falush D, van Dorp L, Lawson D (2016) A tutorial on how (not) to over-interpret STRUCTURE/ADMIXTURE bar plots. bioRxiv, 066431.

Fay RR, Popper AN (2000) Evolution of hearing in vertebrates: the inner ears and processing. Hearing research 149, 1–10.

Fitzpatrick S, Gerberich J, Kronenberger J, Angeloni L, Funk W (2015) Locally adapted traits maintained in the face of high gene flow. Ecology Letters 18, 37–47.

Fitzpatrick SW, Gerberich JC, Angeloni LM, et al. (2016) Gene flow from an adaptively divergent source causes rescue through genetic and demographic factors in two wild populations of Trinidadian guppies. Evolutionary applications 9, 879–891.

Frankham R (2015) Genetic rescue of small inbred populations: metalJanalysis reveals large and consistent benefits of gene flow. Molecular ecology 24, 2610–2618.

Fumey J, Hinaux H, Noirot C, et al. (2018) Evidence for late Pleistocene origin of Astyanax mexicanus cavefish. BMC evolutionary biology 18, 43.

Geneva AJ, Muirhead CA, Kingan SB, Garrigan D (2015) A new method to scan genomes for introgression in a secondary contact model. PloS one 10, e0118621.

Gompel N, PruD’;homme B (2009) The causes of repeated genetic evolution. Developmental Biology 332, 36–47.

Gross J, Wilkens H (2013) Albinism in phylogenetically and geographically distinct populations of Astyanax cavefish arises through the same loss-of-function Oca2 allele. Heredity 111, 122–130.

Gross JB (2012) The complex origin of Astyanax cavefish. BMC evolutionary biology 12, 105.

Gross JB, Borowsky R, Tabin CJ (2009) A novel role for Mc1r in the parallel evolution of depigmentation in independent populations of the cavefish Astyanax mexicanus. PLoS genetics 5, e1000326–e1000326.

Gutenkunst RN, Hernandez RD, Williamson SH, Bustamante CD (2009) Inferring the joint demographic history of multiple populations from multidimensional SNP frequency data. PLoS genetics 5, e1000695.

Hausdorf B, Wilkens H, Strecker U (2011) Population genetic patterns revealed by microsatellite data challenge the mitochondrial DNA based taxonomy of Astyanax in Mexico (Characidae, Teleostei). Molecular Phylogenetics and Evolution 60, 89–97.

Hellenthal G, Busby GB, Band G, et al. (2014) A genetic atlas of human admixture history. Science 343, 747–751.

Hoegg S, Brinkmann H, Taylor JS, Meyer A (2004) Phylogenetic timing of the fish-specific genome duplication correlates with the diversification of teleost fish. Journal of Molecular Evolution 59, 190–203.

Hudson RR (1990) Gene genealogies and the coalescent process. Oxford surveys in evolutionary biology 7, 44.

Hudson RR, Kreitman M, Aguadé M (1987) A test of neutral molecular evolution based on nucleotide data. Genetics 116, 153–153.

Jaggard JB, Robinson BG, Stahl BA, et al. (2017) The lateral line confers evolutionarily derived sleep loss in the Mexican cavefish. Journal of Experimental Biology in press.

Jaggard JB, Stahl BA, Lloyd E, et al. (2018) Hypocretin underlies the evolution of sleep loss in the Mexican cavefish. eLife 7.

Jakobsson M, Edge MD, Rosenberg NA (2013) The relationship between FST and the frequency of the most frequent allele. Genetics 193, 515–528.

Jeffery WR (2001) Cavefish as a model system in evolutionary developmental biology. Developmental Biology 231, 1–12.

Jeffery WR (2009) Chapter 8: Evolution and development in the cavefish Astyanax. Current Topics in Developmental Biology 86, 191–221.

Keene A, Yoshizawa M, McGaugh SE (2015) Biology and Evolution of the Mexican Cavefish Elsevier: Academic Press.

Kowalko JE, Rohner N, Linden TA, et al. (2013a) Convergence in feeding posture occurs through different genetic loci in independently evolved cave populations of Astyanax mexicanus. Proceedings of the National Academy of Sciences USA 110, 16933–16938.

Kowalko JE, Rohner N, Rompani SB, et al. (2013b) Loss of schooling behavior in cavefish through sight-dependent and sight-independent mechanisms. Current Biology 23, 1874–1883.

Krishnan J, Rohner N (2017) Cavefish and the basis for eye loss. Philosophical Transactions Royal Society B 372, 20150487.

Kronenberger J, Funk W, Smith J, et al. (2017) Testing the demographic effects of divergent immigrants on small populations of Trinidadian guppies. Animal Conservation 20, 3–11.

Langecker TG, Wilkens H, Junge P (1991) Introgressive hybridization in the Pachon Cave population of Astyanax fasciatus (Teleostei: Characidae). Ichthyological exploration of freshwaters. Munchen 2, 209–212.

Li H (2013) Aligning sequence reads, clone sequences and assembly contigs with BWA-MEM. arXiv preprint arXiv:1303.3997.

Li H, Durbin R (2009) Fast and accurate short read alignment with Burrows–Wheeler transform. Bioinformatics 25, 1754–1754.

Li H, Durbin R (2010) Fast and accurate long-read alignment with Burrows-Wheeler transform. Bioinformatics 26, 589–589.

Lohse M, Bolger A, Nagel A, et al. (2012) RobiNA: a user-friendly, integrated software solution for RNA-Seq-based transcriptomics. Nucleic Acids Research 40(Web Server issue), W622–627.

Losos JB (2011) Convergence, adaptation, and constraint. Evolution 65, 1827–1840.

Lowry DB, Hoban S, Kelley JL, et al. (2016) Breaking RAD: An evaluation of the utility of restriction site associated DNA sequencing for genome scans of adaptation. Molecular Ecology Resources 17, 142–152.

Malinsky M, Svardal H, Tyers AM, et al. (2017) Whole genome sequences of Malawi cichlids reveal multiple radiations interconnected by gene flow. bioRxiv, 143859.

Martin M (2011) Cutadapt removes adapter sequences from high-throughput sequencing reads. EMBnet.journal 17, 10.

McGaugh SE, Gross JB, Aken B, et al. (2014) The cavefish genome reveals candidate genes for eye loss. Nature communications 5, 5307–5307.

McGaugh SE, Noor MAF (2012) Genomic impacts of chromosomal inversions in parapatric Drosophila species. Philosphilical Transactions of the Royal Society B 367, 422–429.

McKenna A, Hanna M, Banks E, et al. (2010) The Genome Analysis Toolkit: A MapReduce framework for analyzing next-generation DNA sequencing data. Genome research 20, 1297–1303.

Meier JI, Marques DA, Mwaiko S, et al. (2017) Ancient hybridization fuels rapid cichlid fish adaptive radiations. Nature communications 8, 14363.

Meyer A, Van de Peer Y (2005) From 2R to 3R: Evidence for a fishlJspecific genome duplication (FSGD). Bioessays 27, 937–945.

Mitchell RW, Russell WH, Elliott WR (1977) Mexican eyeless characin fishes, genus Astyanax: Environment, distribution, and evolution Texas Tech Press, Texas.

Moran D, Softley R, Warrant EJ (2014) Eyeless Mexican cavefish save energy by eliminating the circadian rhythm in metabolism. PloS one 9, e107877–e107877.

Nair S, Williams JT, Brockman A, et al. (2003) A selective sweep driven by pyrimethamine treatment in southeast asian malaria parasites. Molecular biology and evolution 20, 1526–1536.

Nei M (1987) Molecular Evolutionary Genetics Columbia University Press, New York.

Noor MAF, Bennett SM (2009) Islands of speciation or mirages in the desert? Examining the role of restricted recombination in maintaining species. Heredity 103, 439–444.

O’;Quin KE, Yoshizawa M, Doshi P, Jeffery WR (2013) Quantitative genetic analysis of retinal degeneration in the blind cavefish Astyanax mexicanus. PloS one 8, e57281.

O’Quin K, McGaugh SE (2015) The genetic bases of troglomorphy in Astyanax: How far we have come and where do we go from here? In: Biology and Evolution of the Mexican Cavefish (ed. A. Keene My, S.E. McGaugh). Elsevier.

O’Quin KE, Doshi P, Lyon A, et al. (2015) Complex evolutionary and genetic patterns characterize the loss of scleral ossification in the blind cavefish Astyanax mexicanus. PloS one 10, e0142208.

Ornelas-García CP, Domínguez-Domínguez O, Doadrio I (2008) Evolutionary history of the fish genus Astyanax Baird & Girard (1854)(Actinopterygii, Characidae) in Mesoamerica reveals multiple morphological homoplasies. BMC evolutionary biology 8, 340.

Ornelas-García CP, Pedraza-Lara C (2015) Phylogeny and evolutionary history of A. mexicanus In: Biology and Evolution of the Mexican Cavefish (ed. Keene AY, M. McGaugh, S.E.). Elsevier, San Diego.

Page LM, Burr BM (2011) Peterson field guide to freshwater fishes of North America north of Mexico Houghton Mifflin Harcourt, Boston, Massachusetts, USA.

Panaram K, Borowsky R (2005) Gene flow and genetic variability in cave and surface populations of the Mexican tetra, Astyanax mexicanus (Teleostei: Characidae). Copeia 2005, 409–416.

Paradis E, Claude J, Strimmer K (2004) APE: analyses of phylogenetics and evolution in R language. Bioinformatics 20, 289–290.

Patterson N, Moorjani P, Luo Y, et al. (2012) Ancient admixture in human history. Genetics 192, 1065–1093.

Payseur BA, Rieseberg LH (2016) A genomic perspective on hybridization and speciation. Molecular ecology 25, 2337–2360.

Pease J, Rosenzweig B (2015) Encoding data using biological principles: the Multisample Variant Format for phylogenomics and population genomics. IEEE/ACM transactions on computational biology and bioinformatics/IEEE, ACM.

Pease JB, Hahn MW (2015) Detection and polarization of introgression in a five-taxon phylogeny. Systematic biology 64, 651–662.

Peter BM (2016) Admixture, population structure, and F-statistics. Genetics 202, 1485–1501.

Pickrell JK, Pritchard JK (2012) Inference of population splits and mixtures from genome-wide allele frequency data. PLoS genetics 8, e1002967.

Porter ML, Dittmar K, Pérez-Losada M (2007) How long does evolution of the troglomorphic form take? Estimating divergence times in Astyanax mexicanus. Acta Carsologica 36.

Protas M, Conrad M, Gross JB, Tabin C, Borowsky R (2007) Regressive evolution in the Mexican cave tetra, Astyanax mexicanus. Current Biology 17, 452–454.

Protas M, Hersey C, Kochanek D, et al. (2006) Genetic analysis of cavefish reveals molecular convergence in the evolution of albinism. Nature genetics 38, 107–111.

Protas M, Jeffery WR (2012) Evolution and development in cave animals: from fish to crustaceans. WIREs Developmental Biology 1, 823–845.

Protas M, Tabansky I, Conrad M, et al. (2008) Multi-trait evolution in a cave fish, Astyanax mexicanus. Evolution and Development 10, 196–209.

Reich D, Thangaraj K, Patterson N, Price AL, Singh L (2009) Reconstructing Indian population history. Nature 461, 489–494.

Requena T, Cabrera S, Martín-Sierra C, et al. (2014) Identification of two novel mutations in FAM136A and DTNA genes in autosomal-dominant familial Meniere’;s disease. Human molecular genetics 24, 1119–1126.

Richards EJ, Martin CH (2017) Adaptive introgression from distant Caribbean islands contributed to the diversification of a microendemic adaptive radiation of trophic specialist pupfishes. PLoS genetics 13, e1006919.

Riddle MR, Aspiras AC, Gaudenz K, et al. (2018) Insulin resistance in cavefish as an adaptation to a nutrient-limited environment. Nature 555, 647.

Rieseberg LH, Burke JM (2001) The biological reality of species: gene flow, selection, and collective evolution. Taxon, 47–67.

Ritz KR, Noor MA (2016) Mistaken identity: Another bias in the use of relative genetic divergence measures for detecting interspecies introgression. PloS one 11, e0165032.

Roesti M, Gavrilets S, Hendry AP, Salzburger W, Berner D (2014) The genomic signature of parallel adaptation from shared genetic variation. Molecular ecology 23, 3944–3956.

Rosenblum EB, Parent CE, Brandt EE (2014) The molecular basis of phenotypic convergence. Annual Review of Ecology, Evolution, and Systematics 45, 203–226.

Rougemont Q, Gagnaire PA, Perrier C, et al. (2017) Inferring the demographic history underlying parallel genomic divergence among pairs of parasitic and nonparasitic lamprey ecotypes. Molecular ecology 26, 142–162.

Rougeux C, Bernatchez L, Gagnaire P-A (2017) Modeling the multiple facets of speciation-with-gene-flow toward inferring the divergence history of lake whitefish species pairs (Coregonus clupeaformis). Genome biology and evolution 9, 2057–2074.

Salin K, Voituron Y, Mourin J, Hervant F (2010) Cave colonization without fasting capacities: An example with the fish Astyanax fasciatus mexicanus. Comparative Biochemistry and Physiology Part A: Molecular & Integrative Physiology 156, 451–457.

Schlenke TA, Begun DJ (2004) Strong selective sweep associated with a transposon insertion in Drosophila simulans. Proc Natl Acad Sci U S A 101, 1626–1631.

Slatkin M (2005) Seeing ghosts: the effect of unsampled populations on migration rates estimated for sampled populations. Molecular ecology 14, 67–73.

Stamatakis A (2014) RAxML version 8: a tool for phylogenetic analysis and post-analysis of large phylogenies. Bioinformatics 30, 1312–1313.

Stern DL (2013) The genetic causes of convergent evolution. Nature Reviews Genetics 14, 751.

Stern DL, Orgogozo V (2009) Is genetic evolution predictable? Science 323, 746–751.

Strecker U, Bernatchez L, Wilkens H (2003) Genetic divergence between cave and surface populations of Astyanax in Mexico (Characidae, Teleostei). Molecular ecology 12, 699–710.

Strecker U, Faúndez VH, Wilkens H (2004) Phylogeography of surface and cave Astyanax (Teleostei) from Central and North America based on cytochrome b sequence data. Molecular Phylogenetics and Evolution 33, 469–481.

Strecker U, Hausdorf B, Wilkens H (2012) Parallel speciation in Astyanax cave fish (Teleostei) in Northern Mexico. Molecular Phylogenetics and Evolution 62, 62–70.

Team RC (2014) R: A language and environment for statistical computing. In: R Foundation for Statistical Computing.

Tine M, Kuhl H, Gagnaire P-A, et al. (2014) European sea bass genome and its variation provide insights into adaptation to euryhalinity and speciation. Nature communications 5, 5770.

Van Belleghem S, Vangestel C, De Wolf K, et al. (2018) Evolution at two time frames: polymorphisms from an ancient singular divergence event fuel contemporary parallel evolution. bioRxiv, 255554.

Varatharasan N, Croll RP, FranzlJOdendaal T (2009) Taste bud development and patterning in sighted and blind morphs of Astyanax mexicanus. Developmental Dynamics 238, 3056–3064.

Wagenmakers E-J, Farrell S (2004) AIC model selection using Akaike weights. Psychonomic bulletin & review 11, 192–196.

Welch JJ, Jiggins CD (2014) Standing and flowing: the complex origins of adaptive variation. Molecular ecology 23, 3935–3937.

Whiteley AR, Fitzpatrick SW, Funk WC, Tallmon DA (2015) Genetic rescue to the rescue. Trends in Ecology & Evolution 30, 42–49.

Wilkens H (1971) Genetic interpretation of regressive evolutionary processes: studies on hybrid eyes of two Astyanax cave populations (Characidae, Pisces). Evolution 25, 530–544.

Wilkens H, Strecker U (2003) Convergent evolution of the cavefish Astyanax (Characidae, Teleostei): genetic evidence from reduced eye size and pigmentation. Biological Journal of the Linnean Society 80, 545–554.

Wilkens H, Strecker U (2017) Evolution in the Dark Springer.

Wootton JC, Feng X, Ferdig MT, et al. (2002) Genetic diversity and chloroquine selective sweeps in Plasmodium falciparum. Nature 418, 320.

Yamamoto Y, Byerly MS, Jackman WR, Jeffery WR (2009) Pleiotropic functions of embryonic sonic hedgehog expression link jaw and taste bud amplification with eye loss during cavefish evolution. Developmental Biology 330, 200–211.

Yang Z, Rannala B (2012) Molecular phylogenetics: principles and practice. Nature Reviews Genetics 13, 303–314.

Yoshizawa M, Ashida G, Jeffery DE (2012a) Parental genetic effects in a cavefish adaptive behavior explain disparity between nuclear and mitochondrial DNA. Evolution 66, 2975–2982.

Yoshizawa M, Gorički Š, Soares D, Jeffery WR (2010) Evolution of a behavioral shift mediated by superficial neuromasts helps cavefish find food in darkness. Current Biology 20, 1631–1636.

Yoshizawa M, Robinson B, Duboue ER, et al. (2015) Distinct genetic architecture underlies the emergence of sleep loss and prey-seeking behavior in the Mexican cavefish. BMC Biology 13, 1.

Yoshizawa M, Yamamoto Y, O’;Quin KE, Jeffery WR (2012b) Evolution of an adaptive behavior and its sensory receptors promotes eye regression in blind cavefish. BMC Biology 10, 108.

